# Precise genomic deletions using paired prime editing

**DOI:** 10.1101/2020.12.30.424891

**Authors:** Junhong Choi, Wei Chen, Chase C. Suiter, Choli Lee, Florence M. Chardon, Wei Yang, Anh Leith, Riza M. Daza, Beth Martin, Jay Shendure

**Author notes:** Correspondence to (J.C.), (J.S.). These authors contributed equally (Junhong Choi, Wei Chen).

## Abstract

Technologies that precisely delete genomic sequences in a programmed fashion can be used to study function as well as potentially for gene therapy. The leading contemporary method for programmed deletion uses CRISPR/Cas9 and pairs of guide RNAs (gRNAs) to generate two nearby double-strand breaks, which is often followed by deletion of the intervening sequence during DNA repair. However, this approach can be inefficient and imprecise, with errors including small indels at the two target sites as well as unintended large deletions and more complex rearrangements. Here we describe a prime editing-based method that we term *PRIME-Del*, which induces a deletion using a pair of prime editing gRNAs (pegRNAs) that target opposite DNA strands, effectively programming not only the sites that are nicked but also the outcome of the repair. We demonstrate that *PRIME-Del* achieves markedly higher precision in programming deletions than CRISPR/Cas9 and gRNA pairs. We also show that *PRIME-Del* can be used to couple genomic deletions with short insertions, enabling deletions whose junctions do not fall at protospacer-adjacent motif (PAM) sites. Finally, we demonstrate that lengthening the time window of expression of prime editing components can substantially enhance efficiency without compromising precision. We anticipate that *PRIME-Del* will be broadly useful in enabling precise, flexible programming of genomic deletions, including in-frame deletions, as well as for epitope tagging and potentially for programming rearrangements.

## Introduction

The ability to precisely manipulate the genome can critically enable investigations of the function of specific genomic sequences, including genes and regulatory elements. Within the past decade, CRISPR/Cas9-based technologies have proven transformative in this regard, allowing precise targeting of a genomic locus, with a quickly expanding repertoire of editing or perturbation modalities^1^. Among these, the precise and unrestricted deletion of specific genomic sequences is particularly important, with critical use cases in both functional genomics and gene therapy.

Currently, the leading method for programming genomic deletions uses a pair of CRISPR guide RNAs (gRNAs) that each target a protospacer-adjacent motif (PAM) sequence, generating a pair of nearby DNA double-strand breaks (DSBs). Upon simultaneous cutting of two sites, cellular DNA damage repair factors often ligate two ends of the genome without the intervening sequence^2^ through non-homologous end joining (NHEJ) (**Figure 1a**). Although powerful, this approach has several limitations: 1) An attempt to induce a deletion, particularly a longer deletion, often results in short insertions or deletions (indels; typically less than 10-bp) near one or both DSBs, with or without the intended deletion^3–5^; 2) Other unintended mutations including large deletions and more complex rearrangements can frequently occur, and go undetected for technical reasons^5–8^; 3) DNA double-stranded breaks are a cytotoxic insult^9^; and 4) The junctions of genomic deletions programmed by this method are limited by the distribution of naturally occurring PAM sites. Notwithstanding these limitations, various studies have employed this strategy to great effect, *e.g*. to investigate the function of genes and regulatory elements^5,10,11^, as well as towards gene therapy^12,13^. However, limited precision, DSB toxicity and the inability to program arbitrary deletions have handicapped the utility of CRISPR/Cas9-induced deletions in functional and therapeutic genomics.

**Figure 1.**
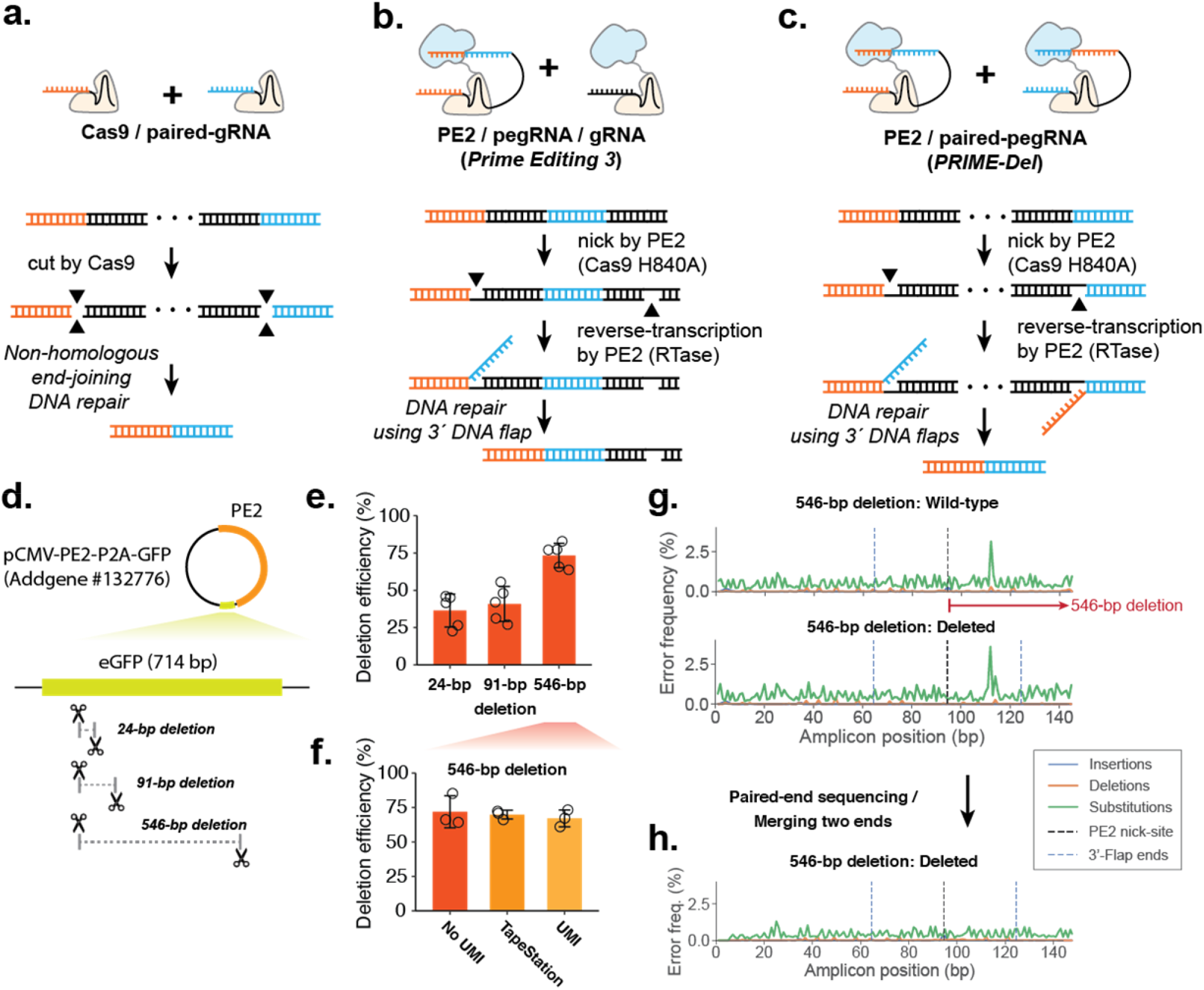
Precise episomal deletions using *PRIME-Del*. **a**. Schematic of Cas9/paired-gRNA deletion strategy. **b**. Schematic of PE3 strategy, wherein the PE2/gRNA complex induces a nick (denoted as a gap in the bottom DNA strand), even after the correct editing event. **c**. Schematic of *PRIME-Del* using pairs of pegRNAs that target opposite DNA strands. Each pegRNA encodes the sites to be nicked at each end of the intended deletion, as well as a 3’ flap that is complementary to the region targeted by the other pegRNA. **d**. Cartoon representation of deletions programmed within the episomally-encoded *eGFP* gene (not drawn to a scale). **e**. *PRIME-Del*-mediated deletion efficiency was measured for 24-bp, 91-bp, and 546-bp deletion experiments. Error bars represent standard deviation for five replicates. **f**. *PRIME-Del*-mediated deletion efficiency was measured for the 546-bp deletion experiment using three methods. Error bars represent standard deviation for three replicates. **g**. Insertion, deletion and substitution error frequencies across sequencing reads from 546-bp deletion experiment. Reads were aligned to reference sequence either without (top) or with (bottom) deletion. Plots are from single-end reads with collapsing of UMIs to reduce sequencing errors; also shown with additional replicates and error-class-specific scales in **Supplementary Fig. 1e**. Note that only one of the two 3’-DNA-flaps is covered by the sequencing read in amplicons lacking the deletion (labeled as ‘wild-type’). **h**. Insertion, deletion and substitution error frequencies across the amplicons from 546-bp deletion experiment after merging paired-end sequencing reads.

Recently, Liu and colleagues described ‘prime editing’, which expands the CRISPR/Cas9 genome editing toolkit in critical ways^14^. Prime-editing utilizes a PE2 enzyme, which is a Cas9 nickase (Cas9 H840A) fused with a reverse-transcriptase, and a 3’-extended gRNA (prime-editing gRNA or pegRNA). The PE2/pegRNA complex can nick one strand of the genome and attach a 3’ single-stranded DNA flap to the nicked site following the template RNA sequence in the pegRNA molecule. By including homologous sequences to the neighboring region, DNA damage repair factors can incorporate the 3’-flap sequence into the genome. The incorporation rate can be further enhanced using an additional gRNA, which makes a nick on the opposite strand, boosting DNA repair with the 3’-flap sequence but often with a decrease in precision (strategy referred to as PE3/PE3b)^14^ (**Figure 1b**). The principal advantage of prime editing lies with its encoding of both the site to be targeted and the nature of the repair within a single molecule, the pegRNA. In addition to demonstrating many other classes of precise edits, Anzalone *et al*. used the PE3 strategy to show that a single pegRNA/gRNA pair could be used to program deletions ranging from 5 to 80 bp achieving high efficiency (52-78%) with modest precision (on average, 11% rate of unintended indels)^14^.

We reasoned that a pair of pegRNAs could be used to specify not only the sites that are nicked but also the outcome of the repair, potentially enabling programming of longer deletions (**Figure 1c**). Here we demonstrate that this strategy, which we call *PRIME-Del*, induces the efficient deletion of sequences up to ∼700 bp in length with much higher precision than observed or expected with either the Cas9/paired-gRNA or PE3 (PE2/pegRNA/gRNA) strategies. We furthermore show that *PRIME-Del* can concurrently program short insertions at the deletion site. Concurrent deletion/insertion can be used to introduce in-frame deletions, to introduce epitope tags concurrently with deletions, and, more generally, to facilitate the programming of deletions unrestricted by the endogenous distribution of PAM sites. By filling these gaps, *PRIME-Del* expands our toolkit to investigate the biological function of genomic sequences at single nucleotide resolution.

## Results

### *PRIME-Del* induces precise deletions in episomal DNA

We first tested the feasibility of the *PRIME-Del* strategy by programming deletions to an episomally encoded *eGFP* gene. We designed pairs of pegRNAs specifying 24-, 91- and 546-bp deletions within the *eGFP* coding region of the pCMV-PE2-P2A-GFP plasmid (Addgene #132776) (**Figure 1d**). We cloned each pair of pegRNAs into a single plasmid with separate promoters, the human U6 and H1 sequences^5^. We transfected HEK293T cells with *eGFP*-targeting paired-pegRNA and pCMV-PE2-P2A-GFP plasmids. We harvested DNA (including both genomic DNA and residual plasmid) from cells 4-5 days after transfection and PCR amplified the *eGFP* region. We then sequenced PCR amplicons to quantify the efficiency of the programmed deletion as well as to detect unintended edits to the targeted sequence.

We calculated deletion efficiency as the number of reads aligning to a reference sequence of the intended deletion, out of the total number of reads aligning to reference sequences either with or without the deletion. Estimated deletion efficiencies ranged from 38% (24-bp deletion) to 77% (546-bp deletion), and were consistent across replicates (note: throughout the paper, the term ‘replicate’ is used to refer to independent transfections) (**Figure 1e**). This result clearly indicates that the *PRIME-Del* strategy outlined in **Fig. 1c** can work. However, we were initially concerned that these were overestimates of efficiency due to the shorter, edited templates being favored by both PCR and Illumina-based sequencing, particularly for the 546-bp deletion, because it has the largest difference between amplicon sizes (766-bp vs. 220-bp for wild-type and deletion amplicons, respectively). To address this, we repeated the amplification on DNA from the 546-bp deletion experiment with a two-step PCR, first adding 15 bp unique molecular identifiers (UMIs) via linear amplification before a second, exponential phase. *PRIME-Del* efficiency was reassessed based on the sequencing data after collapsing of reads with identical UMIs, as well as on the product size distribution (Agilent TapeStation). We observed a slight decrease in deletion efficiency after duplicate removal, from 73% to 66%, comparable to the 70% efficiency measured on the TapeStation (**Figure 1f**). These results suggest that our initial estimates of efficiency are only modestly impacted by size-dependent biases.

For most of these sequencing data, we had only a single read extending over the intended deletion site. As such, it was difficult to distinguish unintended editing outcomes (*e.g*. indels at the nick sites) from PCR or sequencing errors. To address this in part, we plotted frequencies of different classes of errors (substitutions, insertions, deletions) for sequences aligning either to the unedited sequence (**Figure 1g, top**) or the intended deletion (**Figure 1g, bottom**), along the length of the sequencing read. For all replicates of the three deletion experiments (**Supplementary Figure 1**), these profiles showed low rates of substitutions and indels, with nearly identical profiles and no consistent increase in the rate of any class of error at either the positions of the PE2 nick sites or 3’ flap ends above 1%, particularly after collapsing by UMI (**Figure 1g, Supplementary Fig. 1e**) or repeating sequencing with longer, paired-end sequencing reads (**Figure 1h**).

### Simultaneous long deletion and short insertion using *PRIME-Del*

We reasoned that because the homology sequences in the 3’-flaps program the deletion, we could potentially use *PRIME-Del* to concurrently introduce a short insertion at the deletion junction (**Figure 2a**). The desired insertion would be encoded into the pair of pegRNAs in a reverse complementary manner, just 5’ to the deletion-specifying homology sequences. With the conventional strategy for programming deletions, *i.e*. with Cas9 and paired gRNAs, the deletion junctions are determined by the gRNA targets, the selection of which is limited by the natural distribution of PAM sites (**Figure 2b**). Simultaneous long deletion and short insertion with *PRIME-Del* would offer at least three advantages over this conventional strategy. First, an arbitrary insertion of 1-3 bases could enable a reading frame to be maintained after editing, *e.g*. for long deletions intended to remove a protein domain. Second, an arbitrary insertion could be used to effectively move one or both deletion junctions away from the cut-sites determined by the PAM, increasing flexibility to program deletions with base-pair precision. Third, insertion of functional sequences at the deletion junction could allow genome editing with *PRIME-Del* to be coupled to other experimental goals (*e.g*. protein tagging or insertion of a transcriptional start site).

**Figure 2.**
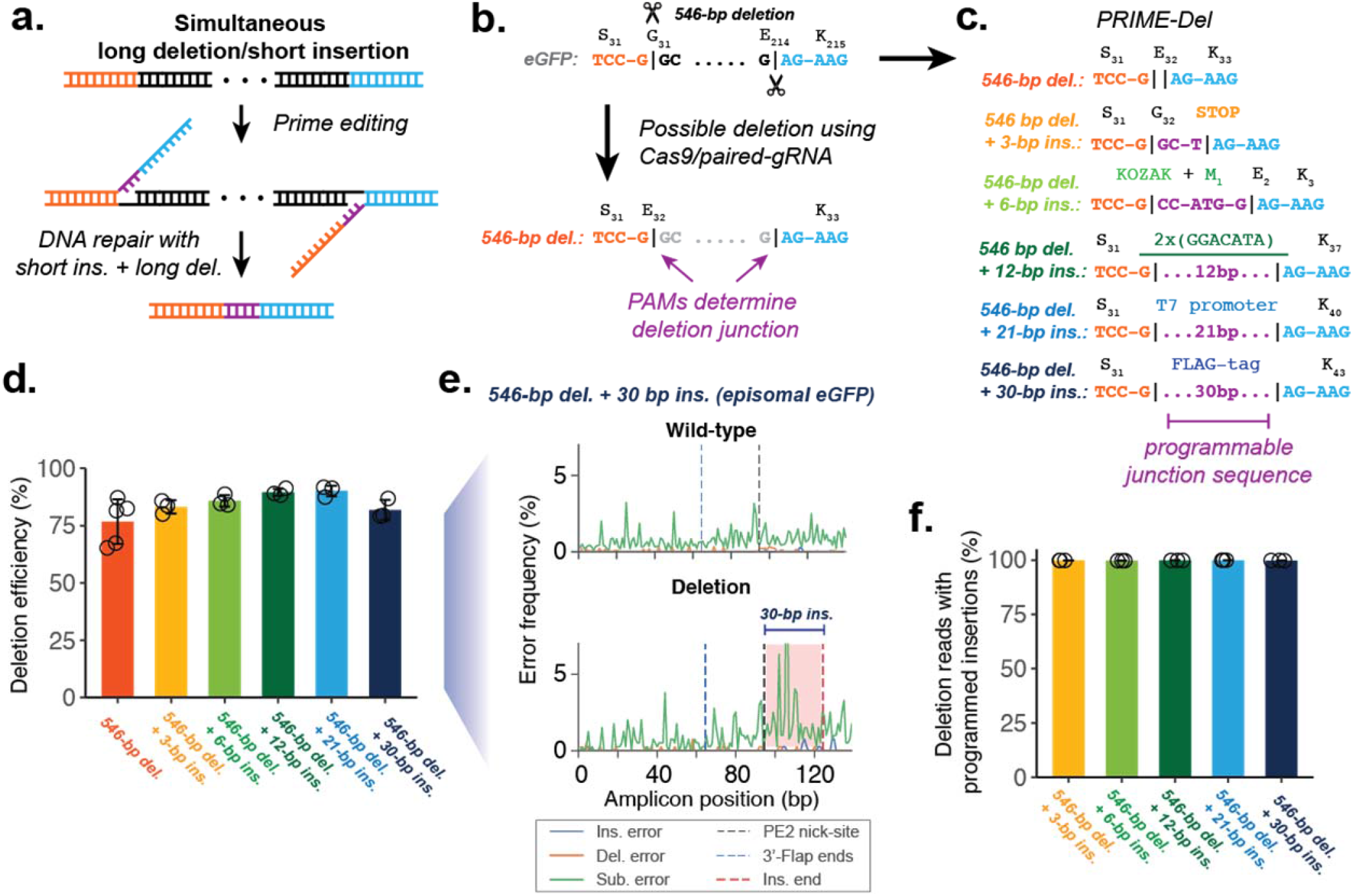
Concurrent programming of deletion and insertion using *PRIME-Del*. **a**. Schematic of strategy, with reverse complementary sequences corresponding to the intended insertion in purple. **b**. Conventional strategy for deletion with Cas9 and pairs of gRNAs. Potential deletion junctions are restricted by the natural distribution of PAM sites. **c**. Pairs of pegRNAs were designed to encode five insertions, ranging in size from 3 to 30 bp, together with a 546 bp deletion in *eGFP*. **d**. Estimated deletion efficiencies in using these pegRNA pairs. Error bars represent standard deviation for at least three replicates. **e**. Representative insertion, deletion and substitution error frequencies plotted across sequencing reads from concurrent 546-bp deletion and 30-bp insertion condition. Plots are from single-end reads without UMI correction. Note that only one of the two 3’-DNA-flaps is covered by the sequencing read in amplicons lacking the deletion (labeled as ‘wild-type’). **f**. The percentage of reads containing the programmed deletion that also contain the programmed insertion. Error bars represent standard deviation for at least three replicates.

To test this concept, we designed pegRNA pairs encoding five insertions ranging from 3 to 30 bp at the junction of a 546-bp programmed deletion within *eGFP* (**Figure 2c**). While our main objective was to test the effect of insertion length on deletion efficiency, we chose insertion sequences for their importance in molecular biology: The 3-bp insertion sequence generates an in-frame stop codon. The 6-bp insertion sequence includes the start codon with the surrounding Kozak consensus sequence. The 12-bp insertion sequence includes tandem repeats of m6A post-transcriptional modification consensus sequence of GGACAT^15^. The 21-bp insertion sequence includes T7 RNA polymerase promoter sequence. The 30-bp insertion sequence encodes for the in-frame FLAG-tag peptide sequence when translated. The estimated efficiencies for simultaneous short insertion and long deletion within the episomal *eGFP* gene were comparable to the 546-bp deletion alone, ranging from 83% to 90% for the various programmed insertions (**Figure 2d**). Also, insertion, deletion and substitution error rates at deletion junctions and across programmed insertions were comparable to the background error frequencies (**Figure 2e, Supplementary Figure 2a**). As expected, the vast majority (>99%) of reads containing the programmed long deletion also contained the insertion (**Figure 2f**), indicating that the full lengths of the pair of 3’-DNA flaps generated following the programmed pegRNA sequences specify the repair outcome (**Figure 2a**).

### *PRIME-Del* induces precise deletions in genomic DNA

Encouraged by our initial results on editing episomal DNA, we next tested *PRIME-Del* on a copy of the *eGFP* gene integrated into the genome. We first generated the polyclonal cell line that carries a single copy of the *eGFP* gene by lentiviral transduction at the multiplicity of infection (MOI) of 0.1, followed by flow-sorting to select GFP-positive cells (**Figure 3a**). We then tested the same pairs of pegRNAs encoding concurrent deletion and insertions (546-bp deletion with or without short insertions at the deletion junction) by transfecting pegRNAs and PE2 without eGFP (pCMV-PE2; Addgene #132775) to these cells. Although editing efficiencies decreased substantially in comparison to episomal *eGFP* (7-17%; **Figure 3b**), we remained unable to detect errors that were clearly associated with editing (**Figure 3c, Supplementary Figure 2b**). Specifically, there was no consistent pattern of error classes above background level accumulating at the nick-site or 3’-DNA-flap incorporation sites. Also, as previously, the vast majority of reads with the 546-bp deletion also contained programmed insertions (**Supplementary Figure 2c**).

**Figure 3.**
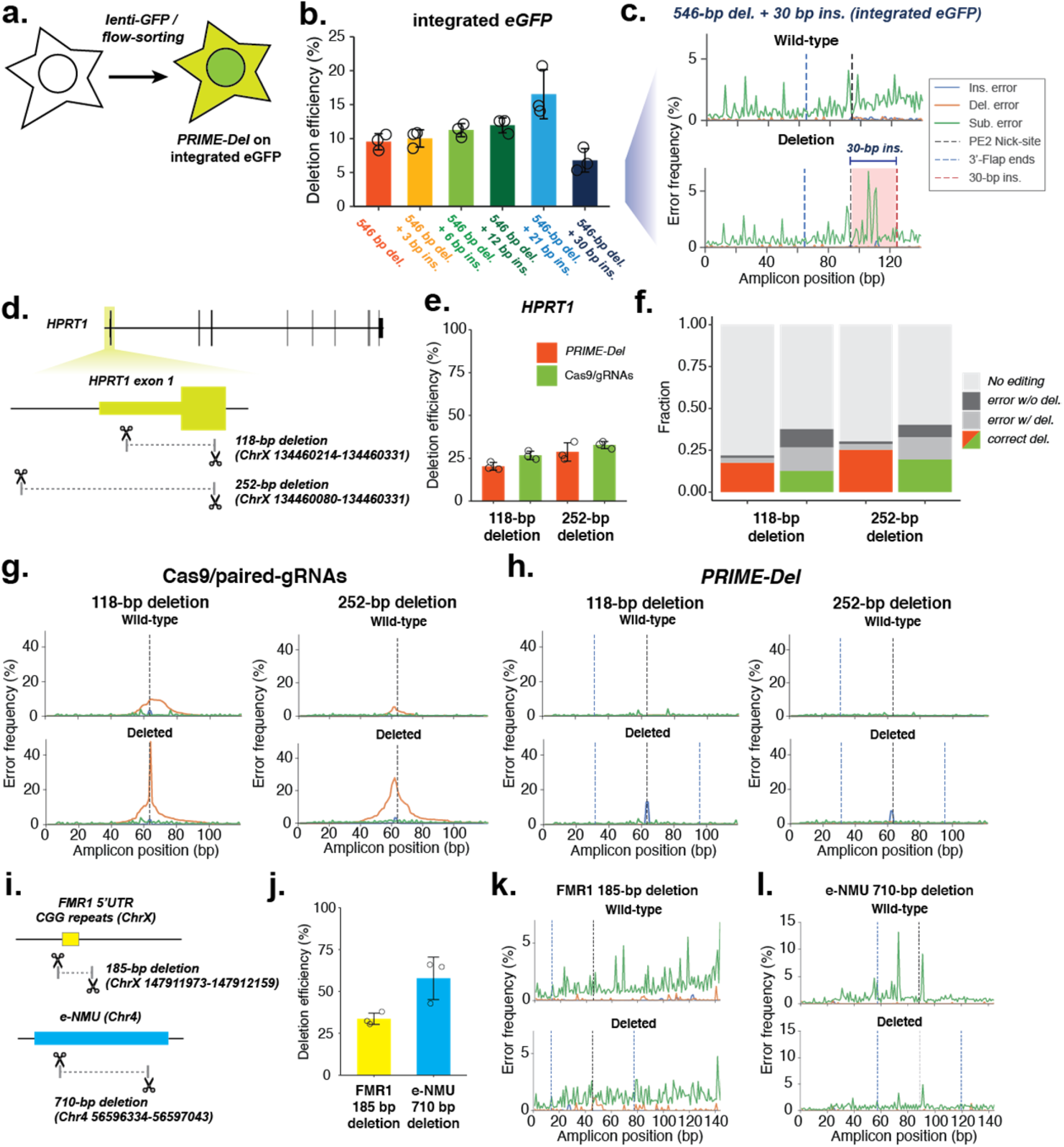
Precise genomic deletions using *PRIME-Del*. **a**. Schematic of generation of the *eGFP*-integrated cell line. **b**. Estimated deletion efficiencies in using *PRIME-Del* for concurrent deletion and insertion on genomically integrated *eGFP*. Error bars represent standard deviation for at least three replicates. **c**. Representative insertion, deletion and substitution error frequencies plotted across sequencing reads from concurrent 546-bp deletion and 30-bp insertion condition on genomically integrated *eGFP*. Plots are from single-end reads without UMI correction. **d**. Cartoon representation of deletions programmed within the *HPRT1* gene. **e**. Deletion efficiencies measured for the 118-bp and 252-bp deletion using either *PRIME-Del* (orange) or Cas9/paired-gRNA (green) strategies. Error bars represent standard deviation for at least three replicates. **f**. Fraction of total reads without indel modifications (“No editing”), indel errors without intended deletion, indel errors with intended deletion, and correct deletion without error. **g**. Representative insertion, deletion and substitution error frequencies plotted across sequencing reads from 118-bp deletion (left) and 252-bp deletion (right) at *HPRT* exon 1, using the Cas9/paired-gRNA strategy. Different error classes are colored the same as in **(c). h**. Same as **(g)**, but for *PRIME-Del* strategy. **i**. Cartoon representation of deletions programmed within the *FMR1* 5’-UTR and an *NMU* enhancer (*“e-NMU*”). **j**. Deletion efficiencies measured for the *FMR1* 5’-UTR (185 bp) and *e-NMU* (710 bp) deletions using *PRIME-Del*. Error bars represent standard deviation for at least three replicates. **k-l**. Representative insertion, deletion and substitution error frequencies plotted across sequencing reads from *PRIME-Del* programmed deletions at *FMR1* (**k**) and *e-NMU* (**l**).

To test *PRIME-Del* on native genes, we designed two pairs of pegRNAs that respectively specified 118 and 252-bp deletions within exon 1 of *HPRT1* (**Figure 3d**). We have previously performed a scanning deletion screen across the *HPRT1* locus using a Cas9/paired-gRNA strategy^5^. To directly compare *PRIME-Del* with Cas9/paired-gRNAs in programming genomic deletions, we also designed two pairs of gRNAs that differ from the corresponding pegRNAs only at their 3’-ends (*i.e*. removing the RT template portion of pegRNA). At exon 1 of *HPRT1*, we observed comparable deletion efficiencies for the *PRIME-Del* and Cas9/paired-gRNA strategies, with nearly 20% and 30% efficiencies for 118-bp and 252-bp deletions, respectively (**Figure 3e**).

As has been shown for other targets^3–5^, the Cas9/paired-gRNA strategy often resulted in errors (mostly short deletions), whether with or without the intended deletion (**Figure 3f; Supplementary Figure 3a**). Of reads lacking the intended 118-bp or 252-bp deletions, 15% or 11% also contained an unintended indel at the observable target site, respectively (these are underestimates, because they only account for one of two target sites) (**Figure 3g, top**). Of reads containing the intended 118-bp or 252-bp deletions, 53% or 40% also contained an unintended indel at the deletion junction, respectively (**Figure 3g, bottom**). Such junctional errors are an established consequence of error-prone repair by NHEJ. In contrast, unintended indels were far less common with *PRIME-Del* (**Figure 3f; Supplementary Figure 3b**). Of reads lacking the intended 118-bp or 252-bp deletions, 1.9% or 2.2% also contained an unintended short indel at the observable target site, respectively (**Figure 3h, top**). Of reads containing the intended 118-bp or 252-bp deletions, 14% or 12% also contained an unintended indel at the deletion junction, respectively (**Figure 3h, bottom**).

For *PRIME-Del*, the observation of an appreciable rate of insertions at the deletion junction in association with intended deletions (**Figure 3h, bottom; Supplementary Figure 3b**) contrasts with our earlier observations at *eGFP*, where these rates were consistently equivalent to background. To explore this further, we performed paired-end sequencing of these amplicons to bidirectionally cover the deletion junction and facilitate removal of PCR duplicates using 15-bp UMI sequences. This revealed that for both pairs of pegRNAs targeting *HPRT1*, these errors corresponded to long insertions (mean 47-bp +/- 12-bp; **Supplementary Figure 4**). The most frequent long insertion at the 118-bp deletion junction was 55-bp, a chimeric sequence between two 32-bp 3’-DNA flap sequences, overlapping at a ‘GCCCT’ sequence, suggesting its origin from the annealing of GC-rich ends of 3’-DNA flaps. Similar chimeric sequences were observed as insertions at the 252-bp deletion junction, overlapping at ‘GCCG’ within their 3’-DNA flaps. Nonetheless, even with these long insertions, 80% and 83% of all reads containing an indel matched the intended deletion exactly with *PRIME-Del*, but only 34% and 49% with the Cas9/paired-gRNA strategies (**Figure 3f**). Indel errors from the Cas9/paired-gRNA strategy are likely underestimated, because errors at only one of two Cas9 cut-sites are captured by our sequencing strategy. Of note, the structure of the observed insertions and the lack of similar errors in applying *PRIME-Del* to the *eGFP* locus (and other loci; see below) suggest that this issue may be addressable through careful pegRNA design.

We further tested genomic deletion using *PRIME-Del* at two additional native loci: the sequence encoding the 5’ untranslated region (5’-UTR) of *FMR1*, and an enhancer of the *NMU* gene (‘*e-NMU*’) (**Figure 3i**). The 5’-UTR of the *FMR1* gene includes a CGG-repeat expansion region that is implicated in Fragile-X syndrome^16^. We designed a pair of pegRNAs encoding a 185-bp deletion at *FMR1* that removes this repeat-expansion region. The *e-NMU* region corresponds to a recently discovered enhancer for *NMU*, experimentally verified using Cas9/paired-gRNA deletions^6,17^. We designed a pair of pegRNAs encoding a 710-bp deletion to delete this enhancer. Using *PRIME-Del*, we observed 34% and 58% deletion efficiencies for the 185-bp *FMR1* and 710-bp *e-NMU* deletions, respectively (**Figure 3j**). In contrast with *HPRT1* exon 1 but similar to *eGFP*, we did not observe recurrent insertions at either of these deletion junctions, further suggesting that this error mode may be specific to certain pegRNA pairs (**Figure 3k**,**l**).

### Extending the editing time window enhances prime editing and *PRIME-Del* efficiency

In contrast with Cas9-mediated DSBs followed by NHEJ, both prime editing and *PRIME-Del* have high editing precision, producing an intended edit or conserving the original editable sequence. We reasoned that if the editing efficiencies of prime editing and *PRIME-Del* are limited by the transient availability of PE2/pegRNA molecules in the cell, extending PE2/pegRNA expression through stable genomic integration or, alternatively, repetitive transfection, would boost the rates of successful editing over time, particularly if uneditable “dead ends” outcomes are not concurrently accruing.

To allow the prolonged expression of PE2 in cells, we generated monoclonal PE2-expressing HEK293T and K562 cell lines (termed HEK293T(PE2) and K562(PE2), respectively). Because the PE2 gene was larger than the lentiviral vector’s typical limit, we cloned PE2 into the piggyBAC cargo, transfected it along with the piggyBAC transposase, and identified a monoclonal cell line with active PE2. To express pegRNAs in the PE2-expressing cell lines, we generated lentiviral vectors with pegRNAs and transduced them into both HEK293T(PE2) and K562(PE2) cells (**Figure 4a**). We tested two different deletions at *HPRT1* using *PRIME-Del* (the aforedescribed 118-bp and 252-bp deletions at exon 1), along with standard prime editing to insert 3-bp (CTT) into the synthetic HEK3 target sequence^14^. In K562(PE2), we observed a steady increase of the correctly edited population over time, both for CTT-insertion using prime editing and for 118- or 252-bp deletions using *PRIME-Del*. The end-point prime editing efficiencies for the CTT-insertion were very high, reaching 90% of targets with correct edits by 19 days after the first transduction of pegRNA into K562(PE2) cells (**Figure 4b**). The rate of precise deletions using *PRIME-Del* also reached nearly 50% and 25% for the 118-bp and 252-bp deletions, respectively, by 19 days. In HEK293T(PE2) cells, we observed lower CTT-insertion efficiencies for the first 10 days, but eventually reaching 80-90% by day 19 (**Figure 4c**). Unexpectedly, we observed the near-absence of *PRIME-Del*-induced deletions in HEK293T(PE2) cells (**Figure 4c**). However, the same HEK293T(PE2) cell line showed modest increases in editing to 5 − 50% when we attempted multiple transfections of either PE2/pegRNA without additional stable integration or PE2 alone after stable integration of piggyBAC-pegRNA, over four weeks (**Supplementary Figure 5**). Together, our results confirm that extended expression of prime editing or *PRIME-Del* components can boost efficiency, but also that *PRIME-Del* is more sensitive to PE2/pegRNA expression level differences between transfection and lentiviral transduction than standard prime editing, presumably because the *PRIME-Del* requires two simultaneous PE2/pegRNA actions.

**Figure 4.**
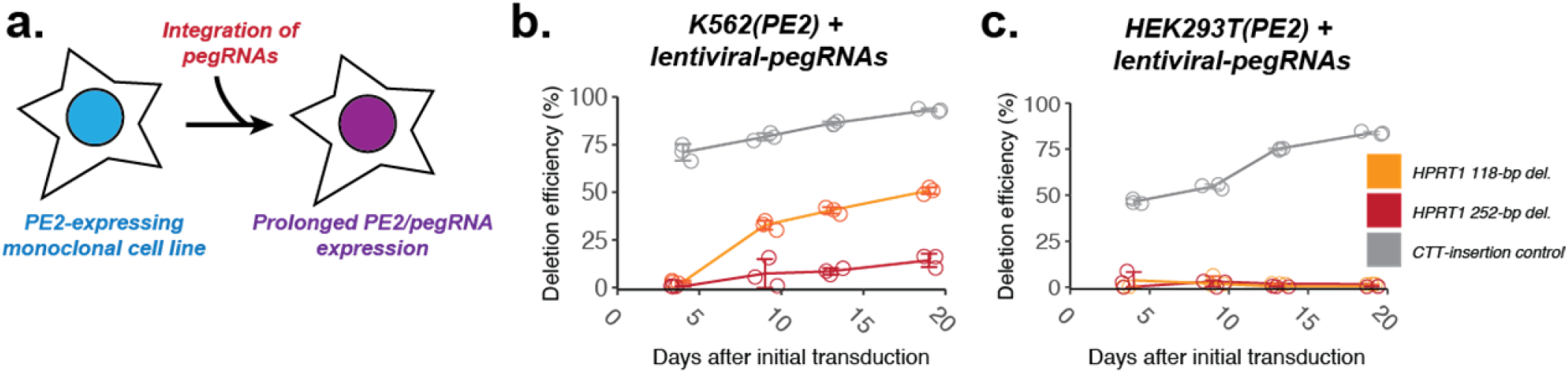
Extending the editing time window enhances prime editing and *PRIME-Del* efficiency. **a**. Schematic for stably expressing both PE2 and pegRNAs via two-step genome integration. **b-c**. Editing efficiencies measured for the 118-bp and 252-bp deletions at genomic *HPRT1* exon 1 using *PRIME-Del* (paired-pegRNA construct) or CTT-insertion using prime editing (single-pegRNA construct) in K562 cells (**b**) or HEK293T cells (**c**), as a function of time after initial transduction of pegRNA(s). Error bars represent standard deviation for three replicates.

## Discussion

Here we introduce *PRIME-Del*, a paired pegRNA strategy for prime editing, and demonstrate that it achieves high precision for programming deletions, both with and without short insertions. We tested deletions ranging from 20 to 700-bp in length at episomal, synthetic genomic, and native genomic loci. The editing efficiency on native genes ranged from 5-50% with a single round of transient transfection in HEK293T cells, although we also observed that prolonged, high expression of prime editing or *PRIME-Del* components enhanced editing efficiency without compromising precision. At the four genomic loci targeted with *PRIME-Del*, we observed high precision of editing except at *HPRT1* exon 1, where long insertions were sometimes observed at the deletion junction (∼13% of edits). Even with these insertion errors, *PRIME-Del* performed better than the conventional Cas9/paired-gRNA strategy, achieving higher efficiencies with fewer errors. Furthermore, the GC-rich ends of 3’-DNA flap sequences of the pegRNA pairs used at *HPRT1* exon 1 appear to underlie the long insertions. Optimizing pegRNA design may be able to eliminate this error mode, as such insertions were not observed in targeting *eGFP*, the 5’-UTR of *FMR1*, or *e-NMU*, experiments for which the pegRNA pairs lacked GC-rich ends at their 3’-DNA flaps. We have developed an accompanying Python-based webtool for designing *PRIME-Del* paired-pegRNA sequences, which notifies the user if such sequences are present in designed pegRNA pairs.

A potential design-related limitation of *PRIME-Del* is that relative to the conventional Cas9/paired-gRNA strategy, it constrains the useable pairs of genomic protospacers, as they need to occur on opposing strands with the PAM sequences oriented towards one another (**Figure 1c**). However, the development and optimization of a near-PAMless^18^ prime editing enzyme would relax this constraint. A further limitation is that because of their longer length, cloning a pair of pegRNAs in tandem is more challenging than cloning gRNA pairs. Each pegRNA used here is 135 to 170 bp in length, such that synthesizing their unique components in tandem as a single, long oligonucleotide approaches the limits of conventional DNA synthesis technology, particularly for goals requiring array-based synthesis of paired pegRNA libraries.

Notwithstanding these limitations, *PRIME-Del* offers significant advantages over alternatives across several potential areas of application (**Figure 5**). Most straightforwardly, *PRIME-Del* can be used for precise programming of long deletions. In addition to the much lower indel error rate observed at the deletion junction compared to the Cas9/paired-gRNA strategy, inducing paired nicks is less likely to result in large, unintended deletions locally, rearrangements genome-wide (chromothripsis), or off-target editing^7,14,19–21^. These characteristics are advantageous for developing therapeutic approaches, *e.g*. where the *PRIME-Del* deletes pathogenic regions such as CGG-repeat expansions in 5’-UTR of *FMR1*, without undesired perturbation of nearby or distant sequences^12,13^.

**Figure 5.**
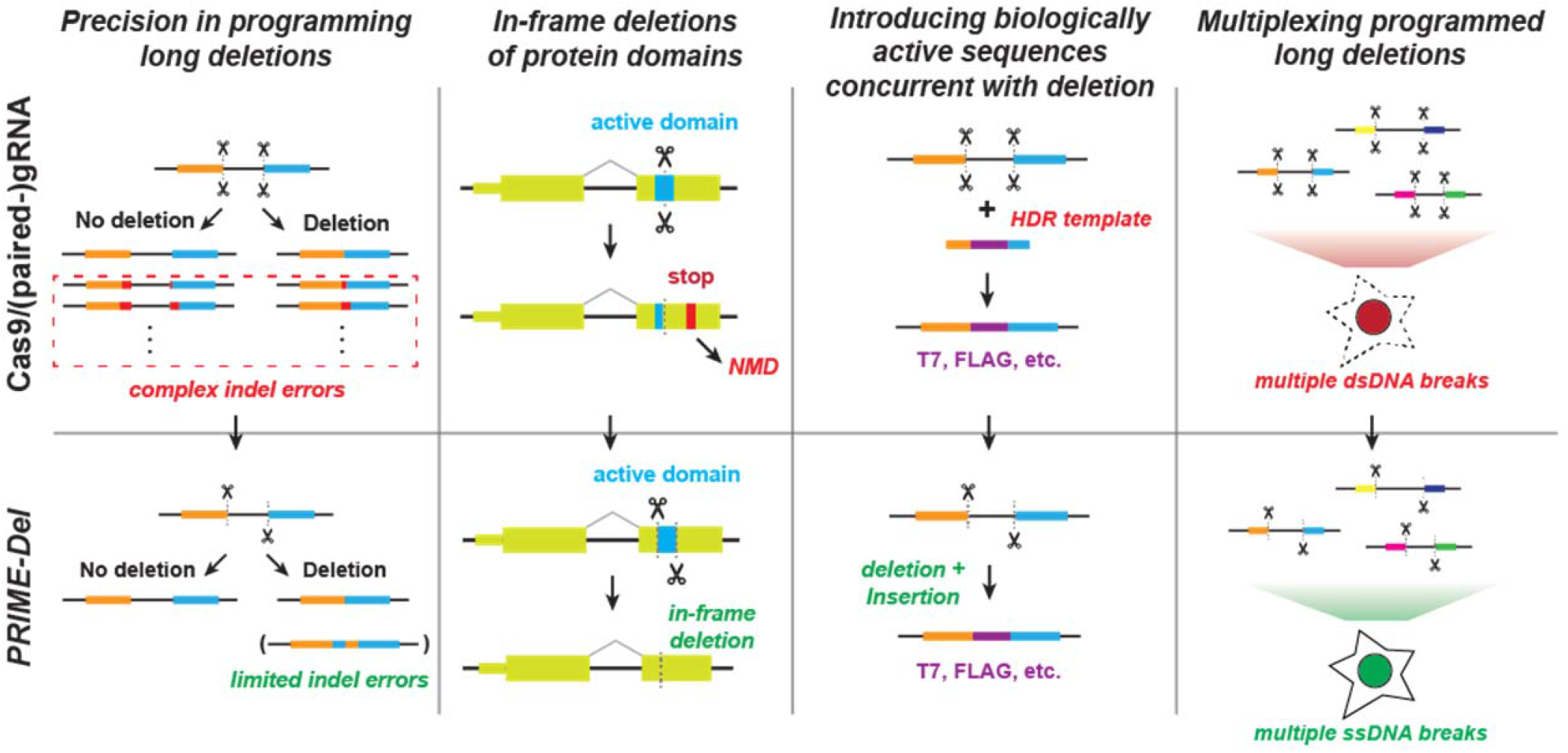
Potential advantages of using *PRIME-Del* in various genome editing applications. The *PRIME-Del* strategy can be used to program precise genomic deletions without generation of short indel errors at Cas9 target sequences. Precision deletion, combined with ability to insert a short arbitrary sequence at the deletion junction, may allow robust gene knockout of active protein domains without generating a premature in-frame stop codon, which can trigger the nonsense-mediated decay (NMD) pathway. *PRIME-Del* also allows replacement of long (<700 bp) genomic regions with arbitrary sequences such as epitope tags or RNA transcription start sites. Single-stranded breaks generated during *PRIME-Del* are likely to be less toxic to the cell, especially when multiple regions are edited in parallel, potentially facilitating its multiplexing.

*PRIME-Del* also allows simultaneous insertion of short sequences at the programmed deletion junction without substantially compromising its efficiency or precision. Inserting short sequences allows for precise deletions of protein domains while preserving the native reading frame, *i.e*. avoiding a premature stop codon that might otherwise elicit a complex nonsense-mediated decay (NMD) response^22,23^. Furthermore, inserting biologically active sequences upon deletion is likely to be advantageous in coupling *PRIME-Del* with technologies, *i.e*. by inserting epitope tags or T7 promoter sequences that can be used as molecular handles within edited genomic loci.

We also expect less toxicity via DNA damage by prime editing-based *PRIME-Del* than with the conventional Cas9/paired-gRNA strategy, which may facilitate multiplexing of programmed genomic deletions for frameworks such as scanDel and crisprQTL^5,6^. For studying the non-coding elements in transcription, efficient and precise deletions up to 700 bp region complements the current use of deactivated Cas9-tethered KRAB domain for CRISPR-interference (CRISPRi), which cannot control the range of epigenetic modification around the target region. As such, we anticipate that *PRIME-Del* could be broadly applied in massively parallel functional assays to characterize native genetic elements at base-pair resolution.

## Supporting information

Supplementary sequences

## Endnotes

## Acknowledgements

We thank former and present members of the Shendure lab including Yi Yin, Jacob Tomes, Silvia Domcke, Alexander Boulgakov, Diego Calderon, Jase Gehring, and Lea Starita for helpful discussions. We thank the David Liu laboratory for sharing the prime editing plasmids. This work was supported by the National Human Genome Research Institute grant 5UM1HG009408-04. J.C. is a Howard Hughes Medical Institute Fellow of the Damon Runyon Cancer Research Foundation (DRG-2403-20). J.S. is an Investigator of the Howard Hughes Medical Institute.

## Author contributions

J.C., C.C.S., and J.S. conceived the project. J.C. designed and performed experiments with guidance from W.C. and J.S. and with assistance from W.C., C.C.S., C.L., F.M.C., A.L., R.M.D., and B.M. F.M.C. and W.Y. contributed to validation data. J.C., W.C., and J.S. analyzed data. W.C. developed the software included in the manuscript. J.C. and J.S. wrote the manuscript with input from other authors.

## Code availability statement

Source code for *PRIME-Del* is available at https://github.com/shendurelab/Prime-del, and interactive webpage for designing pegRNAs for *PRIME-Del* is available at https://primedel.uc.r.appspot.com/.

## Materials and Methods

### pegRNA/gRNA design

For pegRNA/gRNA design, we initially used CRISPOR^24^ to select for 20-bp CRISPR/Cas9 spacers within a given region of interest. We avoided spacers annotated as inefficient, including U6/H1 terminator and GC-rich sequences, and generally selected spacers that had higher predicted efficiencies (Doench scores for U6 transcribed gRNAs^25^). The length of the RT-template portion of a pegRNA was initially set to 30-bp and extended by 1 to 2-bp if it ended in G or C^14,26^.

### Web tool for PRIME-Del paired-pegRNA design

To facilitate *PRIME-Del* paired-pegRNA design, we developed a Python-based web tool that automates the design process. The software takes a FASTA-formatted sequence file as the input, identifies all possible PAM sequences within the provided region, and initially generates all potential paired pegRNA sequences to program deletions. The software can also optionally take as input scored gRNA files generated using Flashfry^27^, CRISPOR^24^ or GPP sgRNA designer^24^; this is highly recommended to identify effective CRISPR/Cas9 spacers. For FlashFry and CRISPOR, gRNA spacers with MIT specificity scores^28^ below 50 are filtered out as recommended by CRISPOR. From initially generated pegRNA pairs, the software selects relevant ones based on additional user-provided design parameters. For example, the user can define the deletion size range. The user can also define the start and end position of desired deletion, and the software will filter to pegRNA pairs present windows centered at those coordinates. pegRNAs for deletions whose junctions do not fall at PAM sites can be designed using the option ‘--precise’ (-p), which adds insertion sequences to both pegRNAs to facilitate the desired edit.

The *PRIME-Del* design software also enables additional design constraints to be specified. The pegRNA RT-template length (also known as the homology arm) is set to 30-bp by default, unless specified otherwise by the user. The pegRNA PBS length is set to 13-bp from the PE2 nick-site by default, unless specified otherwise by the user. The nick position relative to the PAM sequence is predicted using previously identified parameters (Lindel^29^), and RT-template length is adjusted accordingly if the predicted likelihood of generating a nick at a non-canonical position is greater than 25%. PegRNA sequences that include RNA polymerase III terminator sequences (more than four consecutive T’s) are filtered out. The software generates warning messages if more than 4 out of 5 bp in either 3’-DNA-flap are either G or C. Code is available at https://github.com/shendurelab/Prime-del, and interactive webpage is available at https://primedel.uc.r.appspot.com/.

### pegRNA/gRNA cloning

After designing pegRNA/gRNA pairs, we followed the Golden-Gate cloning strategy outlined by Anzalone *et al*.^*14*^, assembling three dsDNA fragments and one plasmid backbone. The first dsDNA fragment contains the pegRNA-1 spacer sequence, annealed from two complementary synthetic single-strand DNA oligonucleotides (IDT) with 4-bp 5’-overhangs. The second dsDNA fragment contains the pegRNA-1 gRNA scaffold sequence, annealed from two DNA oligonucleotides with 5’-end phosphorylation at the end of 4-bp overhang. The third dsDNA fragment contains the pegRNA-1 RT template sequence and primer binding sequence (PBS), pegRNA-1 terminator sequence (six consecutive T’s), and pegRNA-2 sequence with H1 promoter sequence. This was generated by appending pegRNA-1 portion and pegRNA-2 portion to two ends of gene fragments (purchased as gBlocks from IDT) by PCR amplification. The gene fragments contained the pegRNA-1 terminator sequence, H1 promoter sequence, pegRNA-2 spacer sequence, and pegRNA-2 gRNA scaffold sequences. The forward primer included the BsmBI or BsaI restriction site, pegRNA-1 RT template sequence and PBS. The reverse primer included pegRNA-2 RT template, PBS, and BsmBI or BsaI restriction site. PCR fragments (sized between 300 and 400 bp) were purified using 1.0X AMPure (Beckman Coulter) and mixed with two other dsDNA fragments and linearized backbone vector with corresponding overhangs for Golden-Gate-based assembly mix (BsmBI or BsaI golden-gate assembly mix from New England Biolabs). For the pegRNA cloning backbone, we used either the GG-acceptor plasmid (Addgene #132777) or piggyBAC-cargo vector that carries the blasticidin-resistance gene. Each construct plasmid was transformed into Stbl Competent *E. coli* (NEB C3040H) for amplification and purified using a miniprep kit (Qiagen). Cloning was verified using Sanger sequencing (Genewiz).

### Tissue culture, transfection, lentiviral transduction, and monoclonal line generation

HEK293T and K562 cells were purchased from ATCC. HEK293T cells were cultured in Dulbecco’s modified Eagle’s medium with high glucose (GIBCO), supplemented with 10% fetal bovine serum (Rocky Mountain Biologicals) and 1% penicillin-streptomycin (GIBCO). K562 cells were cultured in RPMI 1640 with L-Glutamine (Gibco), supplemented with 10% fetal bovine serum (Rocky Mountain Biologicals) and 1% penicillin-streptomycin (GIBCO). HEK293T and K562 cells were grown with 5% CO2 at 37 C.

For transient transfection, about 50,000 cells were seeded to each well in a 24-well plate and cultured to 70-90% confluency. For prime editing, 375 ng of PE2 plasmid (Addgene #132775) and 125 ng of pegRNA or paired-pegRNA plasmid were mixed and prepared with transfection reagent (Lipofectamine 3000) following the recommended protocol from the vendor. For deletion using Cas9/paired-gRNA, 375 ng of Cas9 plasmid (Addgene #52962) and 125 ng of paired-gRNA plasmid were used instead. Cells were cultured for four to five days after the initial transfection unless noted otherwise, and its genomic DNA was harvested either using DNeasy Blood and Tissue kit (Qiagen) or following cell lysis and protease protocol from Anzalone *et al*.^*14*^.

For lentiviral generation, about 300,000 cells were seeded to each well in a 6-well plate and cultured to 70-90% confluency. Lentiviral plasmid was transfected along with the ViraPower lentiviral expression system (ThermoFisher) following the recommended protocol from the vendor. Lentivirus was harvested following the same protocol, concentrated overnight using Peg-it Virus Precipitation Solution (SBI), and used within 1-2 days to transduce either K562 or HEK293T cells without a freeze-thaw cycle.

For transposase integration, 500 ng of cargo plasmid and 100 ng of Super piggyBAC transposase expression vector (SBI) were mixed and prepared with transfection reagent (Lipofectamine 3000) following the recommended protocol from the vendor. PE2-expressing single-cell clones were generated by integrating PE2 using piggyBAC transposase system, selected by marker (puromycin resistance gene), single-cell sorted into 96-well plates using flow-sort apparatus, cultured for 2-3 weeks until confluency, and screened for PE activity by transfecting CTT-inserting pegRNA alone (Addgene #132778) and sequencing the *HEK3*-target loci.

### DNA sequencing library preparation

To quantify programmed deletion efficiency and possible errors generated by *PRIME-Del*, we amplified the targeted region from purified DNA (∼200 to ∼1000 bp in length) using two-step PCR and sequenced using Illumina sequencing platform (NextSeq or MiSeq) (**Supplementary Figure 1a**). Each purified DNA sample contains wild-type and edited DNA molecules, which were amplified together using the same pairs of primers through each PCR reaction. For the PCR-amplification, we designed a pair of primers for each genomic locus (amplicon) where entire amplicon sizes, with or without deletion, were greater than 200 bp to avoid potential problems in PCR-amplification, in purifying of PCR products, and in clustering onto the sequencing flow-cell.

The first PCR reaction (KAPA Robust) included 300 ng of purified genomic DNA or 2 uL of cell lysate, 0.04 to 0.4 uM of forward and reverse primers in a final reaction volume of 50 uL. Primers included sequencing adapters to their 3’-ends, appending them to both termini of PCR products that amplified genomic DNA. After the first PCR step, products were assessed on 6% TBE-gel and purified using 1.0X AMPure (Beckman Coulter) and added to the second PCR reaction that appended dual sample indexes and flow cell adapters. Products were again purified using AMPure and assessed on the TapeStation (Agilent) before denatured for the sequencing run. For long deletions that generate amplicons sized 200 to 300 bp, we used Miseq sequencing platform at low (8 pM) input DNA concentration to minimize the short amplicons replacing the long amplicons during clustering. Denatured libraries were sequenced using either Illumina NextSeq or MiSeq instruments following the vendor protocols.

For appending 15-bp unique molecular identifiers (UMI), we performed the first PCR reaction in two-steps: First, genomic DNA was linearly amplified in the presence of 0.04 to 0.4 uM of single forward primer in two PCR cycles using KAPA Robust polymerase. This reaction was cleaned up using 1.5X AMPure, and subject to the second PCR with forward and reverse primers. In this case, the forward primer anneals to the upstream of UMI sequence and is not specific to the genomic loci. After PCR amplification, products were cleaned up and added to another PCR reaction that appended dual sample indexes and flow cell adapters, similar to other samples.

### Sequencing data processing and analysis

We designed the sequencing layout to cover at least 50-bp away from the deletion junction in each direction (**Supplementary Figure 1a**). In case of the paired-end sequencing, PEAR^30^ was used to merge the paired-end reads with default parameters and ‘-e’ flag to disable the empirical base frequencies. When 15-bp UMI was present in the sequencing reads, we used a custom Python script to find all reads that share the same UMI, and collapsed into a single read with the most frequent sequence. The resulting sequencing reads were aligned to two reference sequences (with or without deletion) generally using the CRISPResso2 software^31^. Default alignment parameters were used in CRISPResso2, with the gap-open penalty of −20, the gap-extension penalty of −2, and the gap incentive value of 1 for inserting indels at the cut/nick sites. The minimum homology score for a read alignment was explored between 50 and 95 for different amplicon length, and all reported values were generated with a score of 85. Custom python and R scripts were used to analyze the alignment results from CRISPResso2.

Alignment was done using two reference sequences (wild-type and deletion) of same sequence length, generating two sets of reads with respective reference sequences. Deletion efficiencies were calculated as the fraction of total number of reads aligning to the reference sequence with deletion over the total number of reads aligning to either references. Genome editing has three types of error modes: substitution, insertion, and deletion. Each error frequency was plotted across two reference sequences, highlighting in each such plot the Cas9(H840A) nick-site and the 3’-DNA flap incorporation sites.

---

**Supplementary Figure 1.**
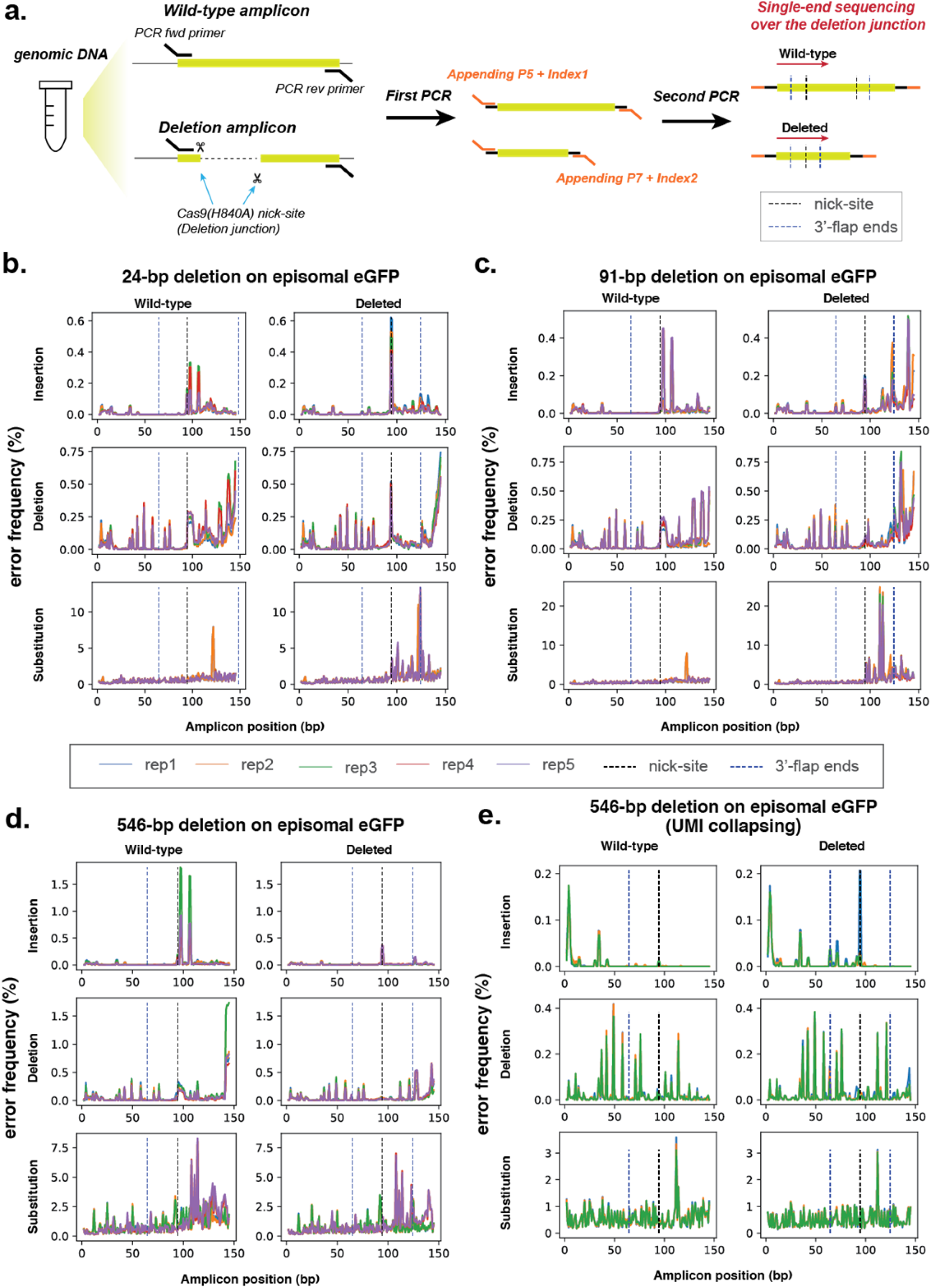
Error profiles with *PRIME-Del* deletions targeting episomally encoded *eGFP*. **a**. Sample preparation schematic for amplicon sequencing. Region around the segment targeted for deletion is amplified from the genomic DNA using two-step PCR amplification that appends sequencing adaptors in the second step. **b-d**. Insertion, deletion and substitution error frequencies across sequencing reads for 24-bp deletion (**b**), 91-bp deletion (**c**), and 546-bp deletion (**d**). These are based on single-end sequencing, with five replicates per experiment, all sequenced on one run, overlaid. Note that except for 24-bp deletion, only one of the two 3’-DNA-flaps is covered by the sequencing read in amplicons lacking the deletion (labeled as ‘wild-type’). *Y*-axis scaling is different for each plot. **e**. Error frequencies across 546-bp deletion after repeating amplification to allow unique molecular identifier (UMI) correction. PCR duplicates identified by UMIs were collapsed into a single read by taking the most frequent sequence sharing the same UMI. These are based on single-end sequencing, with three replicates per experiment, all sequenced on one run, overlaid. *Y*-axis scaling is different for each plot.

**Supplementary Figure 2.**
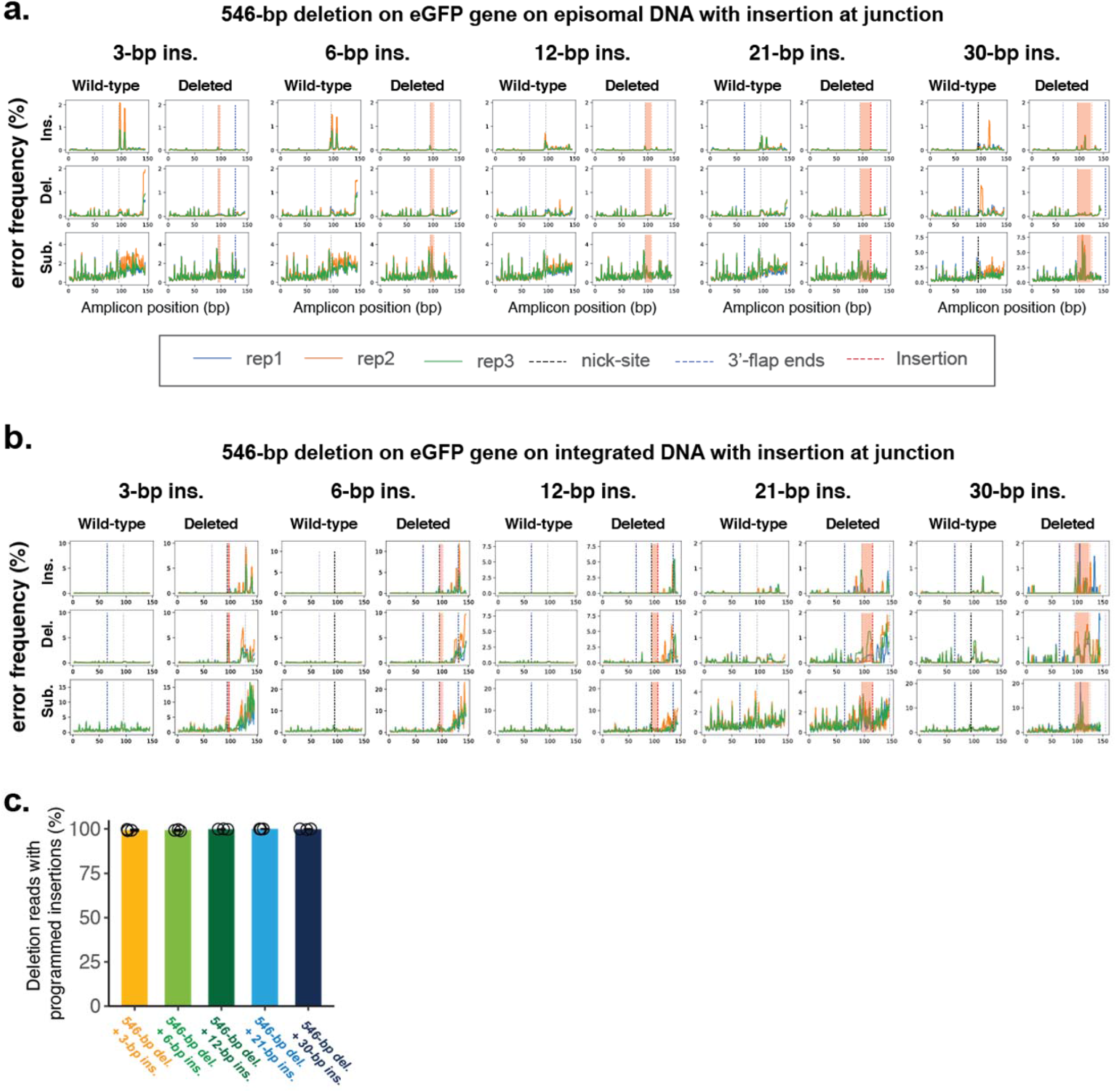
Error profiles with concurrent deletion and insertion at episomally or genomically encoded *eGFP*. **a**. Insertion, deletion and substitution error frequencies plotted across sequencing reads from concurrent 546-bp deletion and various insertion conditions, targeting episomally encoded *eGFP*. These are based on single-end sequencing, with three replicates per experiment, all sequenced on one run, overlaid. Note that only one of the two 3’-DNA-flaps is covered by the sequencing read in amplicons lacking the deletion (labeled as ‘wild-type’). Locations within read corresponding to insertions at deletion junction are highlighted between the nick-site (black dotted line) and end of insertion (red dotted line). *Y*-axis scaling is different for each plot. **b**. Same as (**a**), but for experiments targeting a genomically integrated copy of *eGFP*. **c**. The percentage of reads containing the programmed deletion that also contain the programmed insertion. Similar to **Fig. 2f**, but for experiments targeting a genomically integrated copy of *eGFP*. Error bars represent standard deviation for at least three replicates.

**Supplementary Figure 3.**
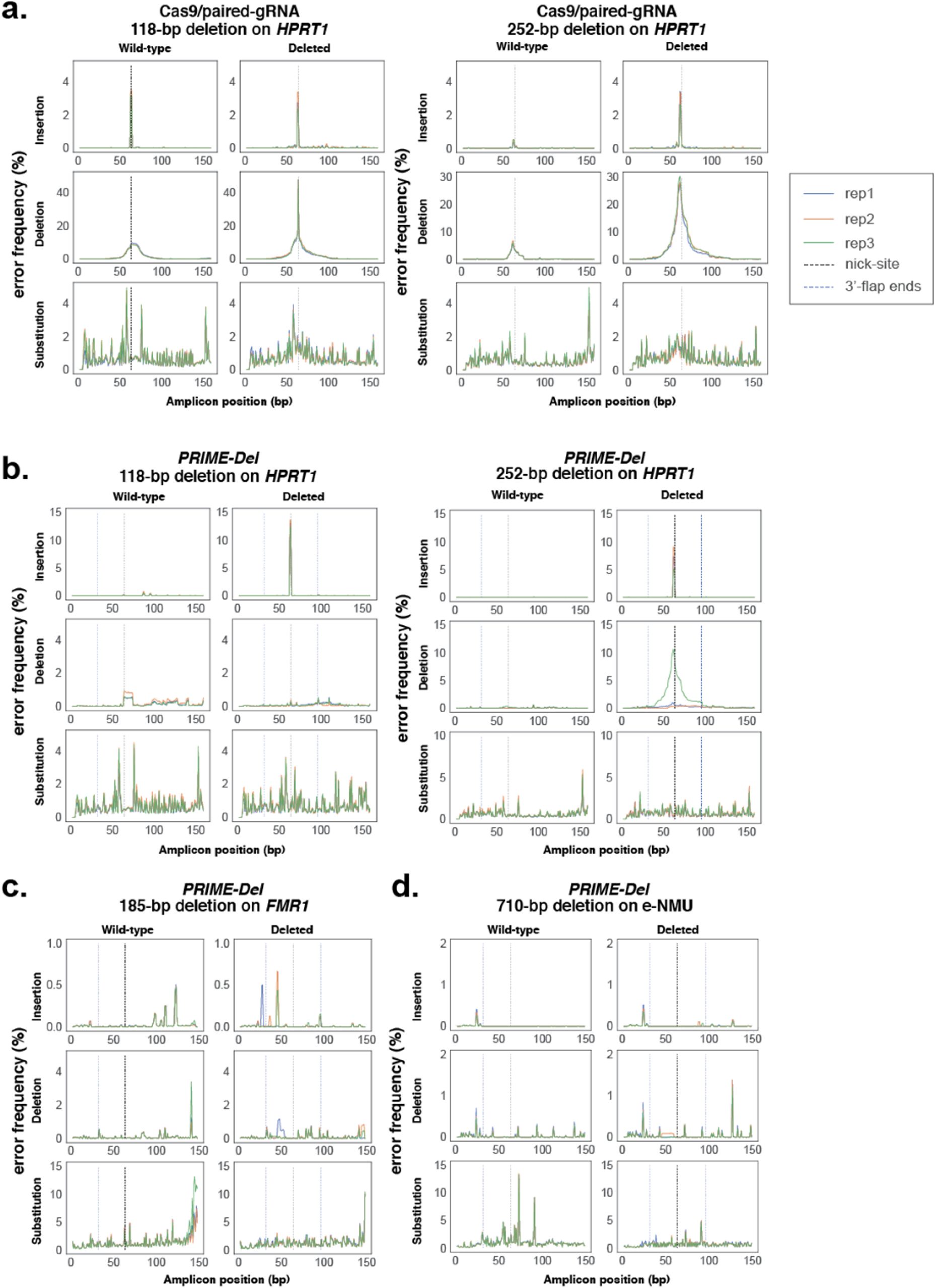
Error profiles with targeting of native *HPRT1, FMR1*, and *e-NMU* loci. **a-c**. Insertion, deletion and substitution error frequencies plotted across sequencing reads from: (**a**) 118-bp or 252-bp deletion on *HPRT1* using the Cas9/paired-gRNA strategy, (**b**) 118-bp or 252-bp deletion on *HPRT1* using the *PRIME-Del* strategy, **(c)** 185-bp deletion on *FMR1* using the *PRIME-Del* strategy, and (**d**) 710-bp deletion on *e-NMU* using the *PRIME-Del* strategy. Sequencing reads aligning to the ‘deletion’ reference for *HPRT1* condition are based on paired-end sequencing, while all the other conditions are based on the single-end sequencing. Each experiment has three replicates sequenced on one run, overlaid. Note that only one of the two 3’-DNA-flaps is covered by the sequencing read in amplicons lacking the deletion (labeled as ‘wild-type’) and that *y*-axis scaling is different for each plot. There may be a sample cross-contamination between Cas9/paired-gRNA replicate #1 and *PRIME-Del* replicate #3, based on the similarity of their deletion error profiles. Those two samples were PCR-amplified and AMPure processed in PCR tubes next to each other.

**Supplementary Figure 4.**
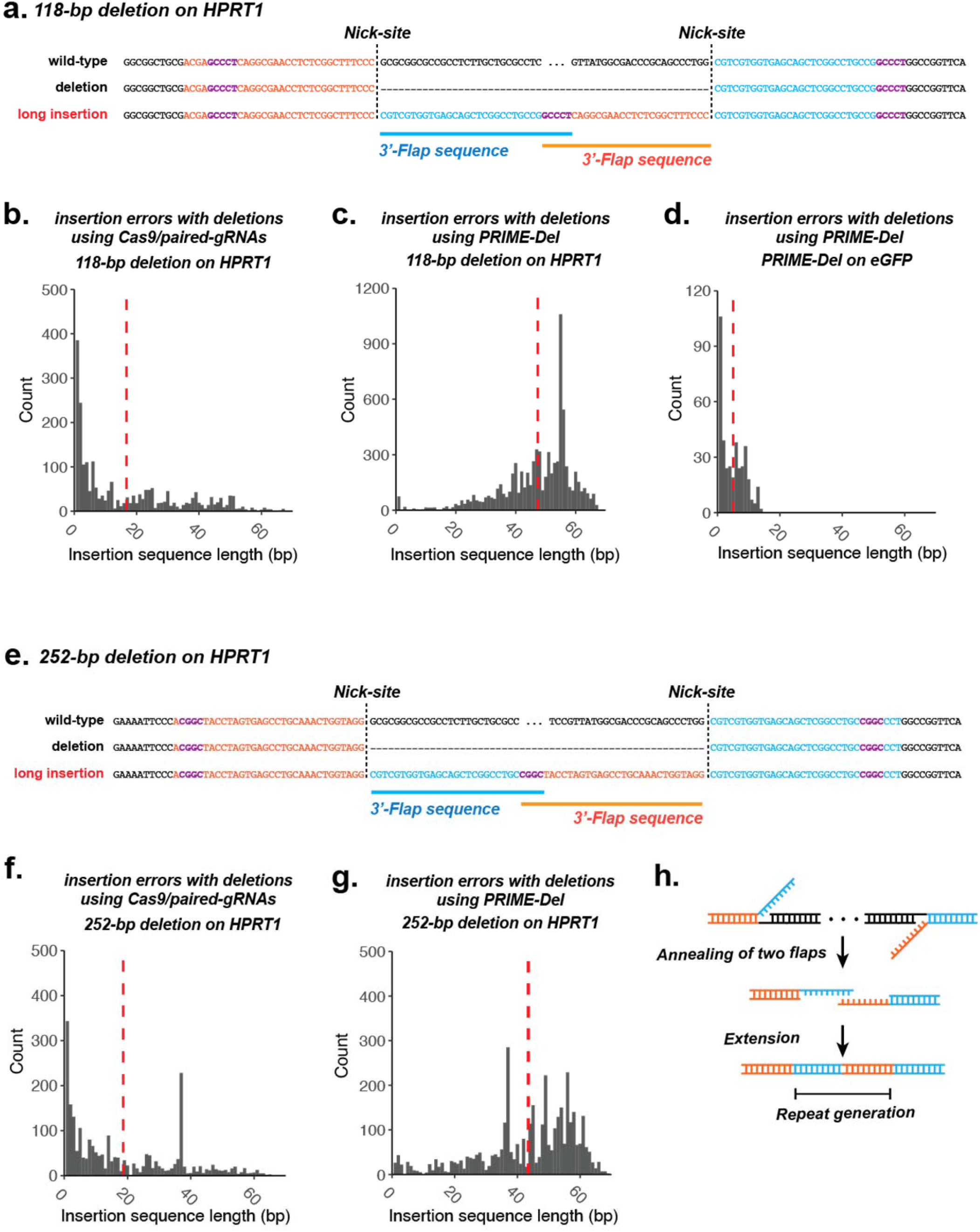
Rare long insertions upon *PRIME-Del* editing of the *HPRT1* exon 1. **a**. We performed paired-end sequencing of amplicons derived from the *PRIME-Del-*edited *HPRT1* locus to bidirectionally cover the deletion junction and facilitate removal of PCR duplicates using 15-bp UMI sequences. This revealed recurrent long insertions that upon inspection appear to be chimeras of the two 3’ flap sequences, with overlap at their GC-rich ends (highlighted in purple). Shown here is a representative insertion from the 118-bp deletion condition. **b-d**. Histograms of insertion sequence lengths for *HPRT1* 118-bp deletion with Cas9/paired-gRNA (**b**), *HPRT1* 118-bp deletion with *PRIME-Del* (**c**), or *eGFP* 546-bp deletion with *PRIME-Del* (**d**). Red vertical lines denote the mean insertion lengths. **e**. Same as (**a**), but representative insertion from the 252-bp deletion condition, also a chimera of the two 3’ flap sequences, with overlap at their GC-rich ends. **f-g**. Histogram of insertion sequence lengths for *HPRT1* 252-bp deletion with *PRIME-Del* (**f**) or Cas9/paired-gRNA (**g**). **h**. Potential mechanism of long insertions with *PRIME-Del*. GC-rich ends of 3’-flaps of paired pegRNAs (*GCCCT* in case of 118-bp deletion and *CGGC* in case of 252-bp deletion) anneal to one another, or to another GC-rich stretch, resulting in insertion upon repair.

**Supplementary Figure 5.**
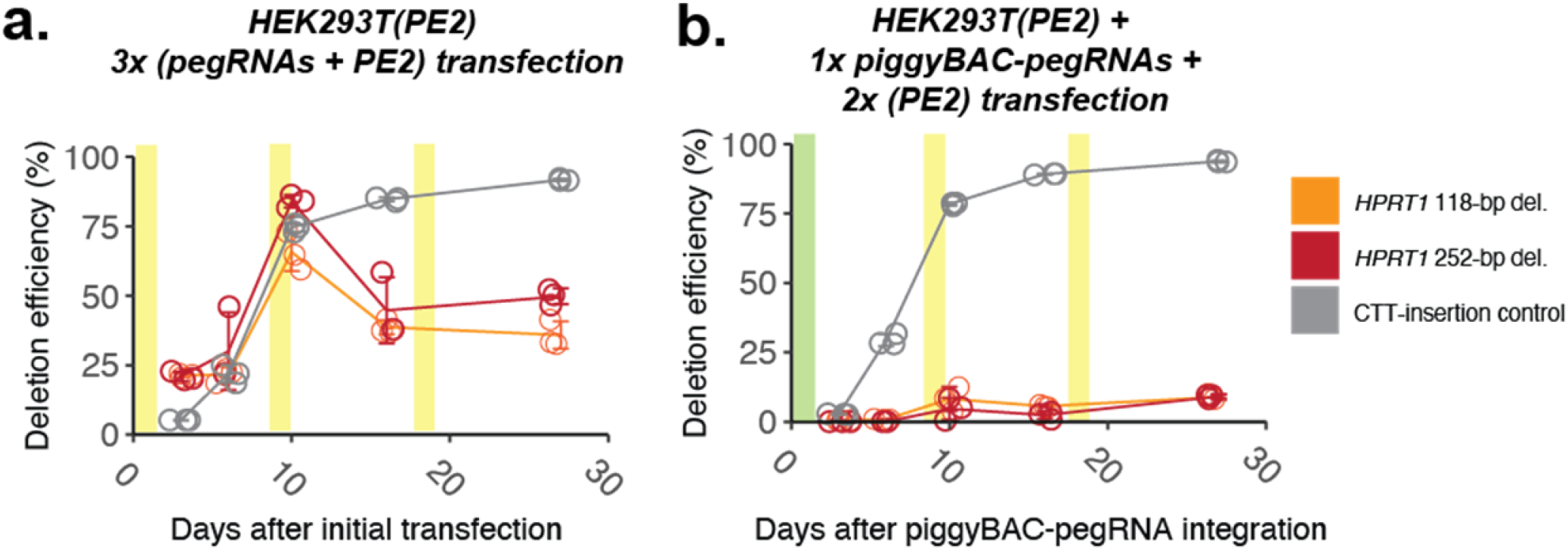
Multiple transfections enhance *PRIME-Del* efficiency in monoclonal HEK293T(PE2) cells. **a**. Editing efficiencies measured for the 118-bp and 252-bp deletions at genomic *HPRT1* exon 1 using *PRIME-Del* (paired-pegRNA construct) or CTT-insertion using prime-editing (single-pegRNA construct), as a function of time after initial transduction of pegRNA(s). Plasmids bearing paired-pegRNAs and PE2 were transfected 3 times (days 0, 9, 18; highlighted in yellow) into PE2-expressing HEK293T cells. Error bars represent standard deviation for three replicates. **b**. Same as (**a**), but first with integration of pegRNAs to PE2-expressing HEK293T via piggyBAC transposon system on Day 0 (highlighted in green), followed by two additional transfections of plasmid bearing PE2 only on Day 9 and 18 (highlighted in yellow). Error bars represent standard deviation for three replicates.

## References

1. Knott, G. J. & Doudna, J. A. CRISPR-Cas guides the future of genetic engineering. Science vol. 361 866–869 (2018).

2. Cong, L. et al. Multiplex genome engineering using CRISPR/Cas systems. Science 339, 819–823 (2013).

3. Canver, M. C. et al. Characterization of genomic deletion efficiency mediated by clustered regularly interspaced short palindromic repeats (CRISPR)/Cas9 nuclease system in mammalian cells. J. Biol. Chem. 289, 21312–21324 (2014).

4. Byrne, S. M., Ortiz, L., Mali, P., Aach, J. & Church G. M. Multi-kilobase homozygous targeted gene replacement in human induced pluripotent stem cells. Nucleic Acids Res. 43, e21 (2015).

5. Gasperini, M. et al. CRISPR/Cas9-Mediated Scanning for Regulatory Elements Required for HPRT1 Expression via Thousands of Large, Programmed Genomic Deletions. Am. J. Hum. Genet. 101, 192–205 (2017).

6. Gasperini, M. et al. A Genome-wide Framework for Mapping Gene Regulation via Cellular Genetic Screens. Cell 176, 1516 (2019).

7. Kosicki, M., Tomberg, K. & Bradley, A. Repair of double-strand breaks induced by CRISPR-Cas9 leads to large deletions and complex rearrangements. Nat. Biotechnol. 36, 765–771 (2018).

8. Zuccaro, M. V. et al. Allele-Specific Chromosome Removal after Cas9 Cleavage in Human Embryos. Cell (2020) doi:10.1016/j.cell.2020.10.025.

9. Mehta, A. & Haber, J. E. Sources of DNA double-strand breaks and models of recombinational DNA repair. Cold Spring Harb. Perspect. Biol. 6, a016428 (2014).

10. Diao, Y. et al. A tiling-deletion-based genetic screen for cis-regulatory element identification in mammalian cells. Nat. Methods 14, 629–635 (2017).

11. Zhu, S. et al. Genome-scale deletion screening of human long non-coding RNAs using a paired-guide RNA CRISPR-Cas9 library. Nat. Biotechnol. 34, 1279–1286 (2016).

12. Khosravi, M. A. et al. Targeted deletion of BCL11A gene by CRISPR-Cas9 system for fetal hemoglobin reactivation: A promising approach for gene therapy of beta thalassemia disease. Eur. J. Pharmacol. 854, 398–405 (2019).

13. Dastidar, S. et al. Efficient CRISPR/Cas9-mediated editing of trinucleotide repeat expansion in myotonic dystrophy patient-derived iPS and myogenic cells. Nucleic Acids Res. 46, 8275–8298 (2018).

14. Anzalone, A. V. et al. Search-and-replace genome editing without double-strand breaks or donor DNA. Nature 576, 149–157 (2019).

15. Dominissini, D. et al. Topology of the human and mouse m6A RNA methylomes revealed by m6A-seq. Nature 485, 201–206 (2012).

16. Verkerk, A. J. et al. Identification of a gene (FMR-1) containing a CGG repeat coincident with a breakpoint cluster region exhibiting length variation in fragile X syndrome. Cell 65, 905–914 (1991).

17. Tippens, N. D. et al. Transcription imparts architecture, function and logic to enhancer units. Nat. Genet. 52, 1067–1075 (2020).

18. Walton, R. T., Christie, K. A., Whittaker, M. N. & Kleinstiver, B.P. Unconstrained genome targeting with near-PAMless engineered CRISPR-Cas9 variants. Science 368, 290–296 (2020).

19. Schene, I. F. et al. Prime editing for functional repair in patient-derived disease models. 2020.06.09.139782 (2020) doi:10.1101/2020.06.09.139782.

20. Owens, D. D. G. et al. Microhomologies are prevalent at Cas9-induced larger deletions. Nucleic Acids Res. 47, 7402–7417 (2019).

21. Kim, D. Y. et al. Unbiased investigation of specificities of prime editing systems in human cells. Nucleic Acids Research (2020) doi:10.1093/nar/gkaa764.

22. El-Brolosy, M. A. et al. Genetic compensation triggered by mutant mRNA degradation. Nature 568, 193– 197 (2019).

23. Ma, Z. et al. PTC-bearing mRNA elicits a genetic compensation response via Upf3a and COMPASS components. Nature 568, 259–263 (2019).

24. Concordet, J.-P. & Haeussler, M. CRISPOR: intuitive guide selection for CRISPR/Cas9 genome editing experiments and screens. Nucleic Acids Res. 46, W242–W245 (2018).

25. Doench, J. G. et al. Optimized sgRNA design to maximize activity and minimize off-target effects of CRISPR-Cas9. Nat. Biotechnol. 34, 184–191 (2016).

26. Kim, H. K. et al. Predicting the efficiency of prime editing guide RNAs in human cells. Nat. Biotechnol. (2020) doi:10.1038/s41587-020-0677-y.

27. McKenna, A. & Shendure, J. FlashFry: a fast and flexible tool for large-scale CRISPR target design. BMC Biol. 16, 74 (2018).

28. Hsu, P. D. et al. DNA targeting specificity of RNA-guided Cas9 nucleases. Nat. Biotechnol. 31, 827–832 (2013).

29. Chen, W. et al. Massively parallel profiling and predictive modeling of the outcomes of CRISPR/Cas9-mediated double-strand break repair. Nucleic Acids Research vol. 47 7989–8003 (2019).

30. Zhang, J., Kobert, K., Flouri, T. & Stamatakis, A. PEAR: a fast and accurate Illumina Paired-End reAd mergeR. Bioinformatics 30, 614–620 (2014).

31. Clement, K. et al. CRISPResso2 provides accurate and rapid genome editing sequence analysis. Nat. Biotechnol. 37, 224–226 (2019).

