## Supplementary Figures for "Precise genomic deletions using paired prime editing"

List of Supplementary Figures:

1. Error profiles with *PRIME-Del* deletions targeting episomally encoded *eGFP*.
2. Error profiles with concurrent deletion and insertion at episomally or genomically encoded *eGFP*.
3. Error profiles with targeting of native *HPRT1*, *FMR1*, and *e-NMU* loci.
4. Rare long insertions upon *PRIME-Del* editing of the *HPRT1* exon 1.
5. Multiple transfections enhance *PRIME-Del* efficiency in monoclonal HEK293T(PE2) cells.

List of Supplementary Data:

1. Sequences of pegRNA and gRNA used in experiments
2. Sequences of primers used for genomic DNA amplification

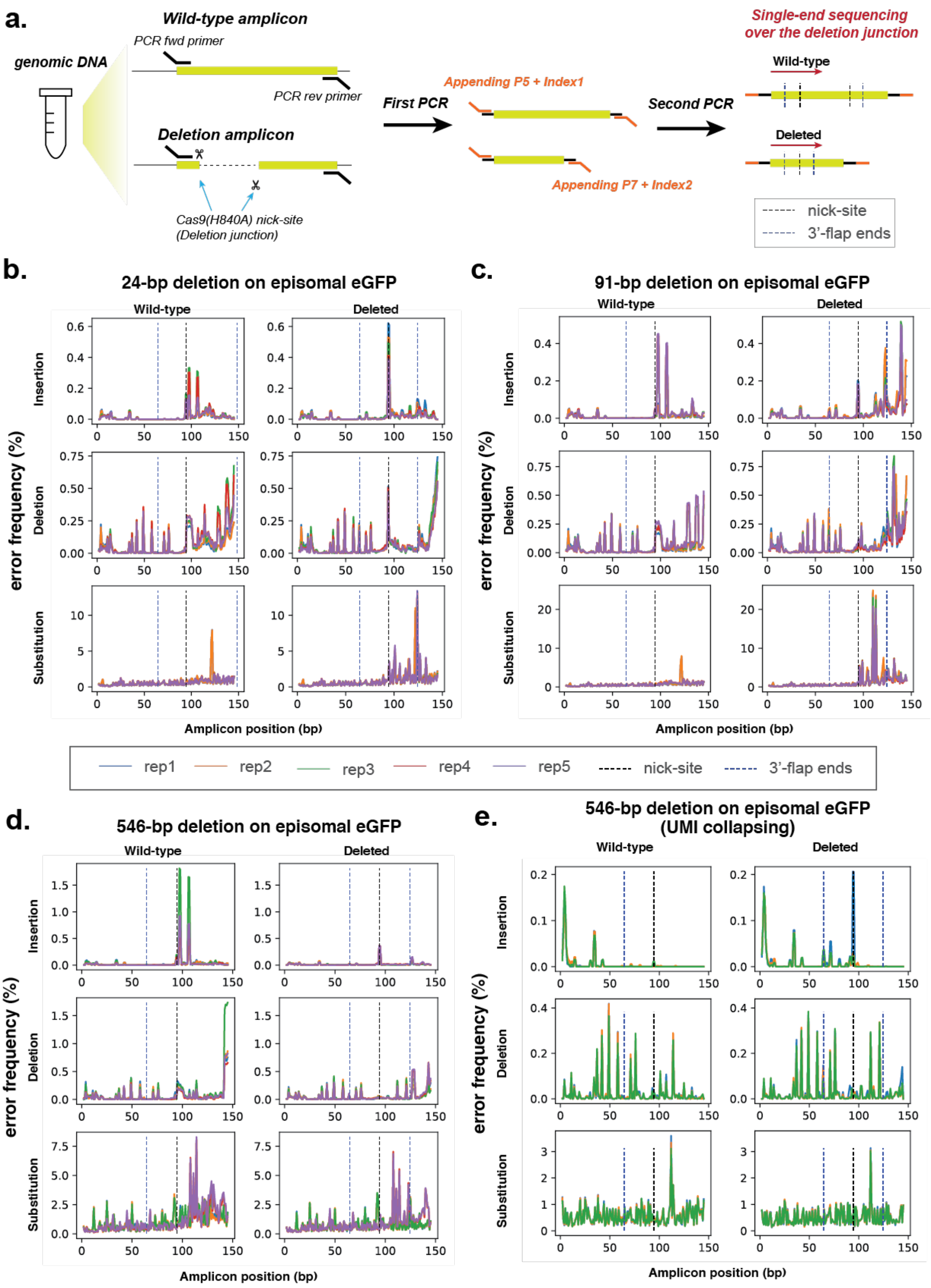
a.
Wild-type amplicon
PCR fwd primer
genomic DNA
PCR rev primer
Deletion amplicon
Cas9(H840A) nick-site
(Deletion junction)
First PCR
Appending P5 + Index1
Appending P7 + Index2
Second PCR
Single-end sequencing
over the deletion junction
Wild-type
Deleted
nick-site
3'-flap ends
b.
24-bp deletion on episomal eGFP
c.
91-bp deletion on episomal eGFP
Wild-type
Deleted
Insertion
Deletion
Substitution
error frequency (%)
Amplicon position (bp)
rep1
rep2
rep3
rep4
rep5
nick-site
3'-flap ends
d.
546-bp deletion on episomal eGFP
e.
546-bp deletion on episomal eGFP
(UMI collapsing)

**Supplementary Figure 1. Error profiles with *PRIME-Del* deletions targeting episomally encoded *eGFP*. a.** Sample preparation schematic for amplicon sequencing. Region around the segment targeted for deletion is amplified from the genomic DNA using two-step PCR amplification that appends sequencing adaptors in the second step. **b-d.** Insertion, deletion and substitution error frequencies across sequencing reads for 24-bp deletion (**b**), 91-bp deletion (**c**), and 546-bp deletion (**d**). These are based on single-end sequencing, with five replicates per experiment, all sequenced on one run, overlaid. Note that except for 24-bp deletion, only one of the two 3'-DNA-flaps is covered by the sequencing read in amplicons lacking the deletion (labeled as 'wild-type'). *Y*-axis scaling is different for each plot. **e.** Error frequencies across 546-bp deletion after repeating amplification to allow unique molecular identifier (UMI) correction. PCR duplicates identified by UMIs were collapsed into a single read by taking the most frequent sequence sharing the same UMI. These are based on single-end sequencing, with three replicates per experiment, all sequenced on one run, overlaid. *Y*-axis scaling is different for each plot.

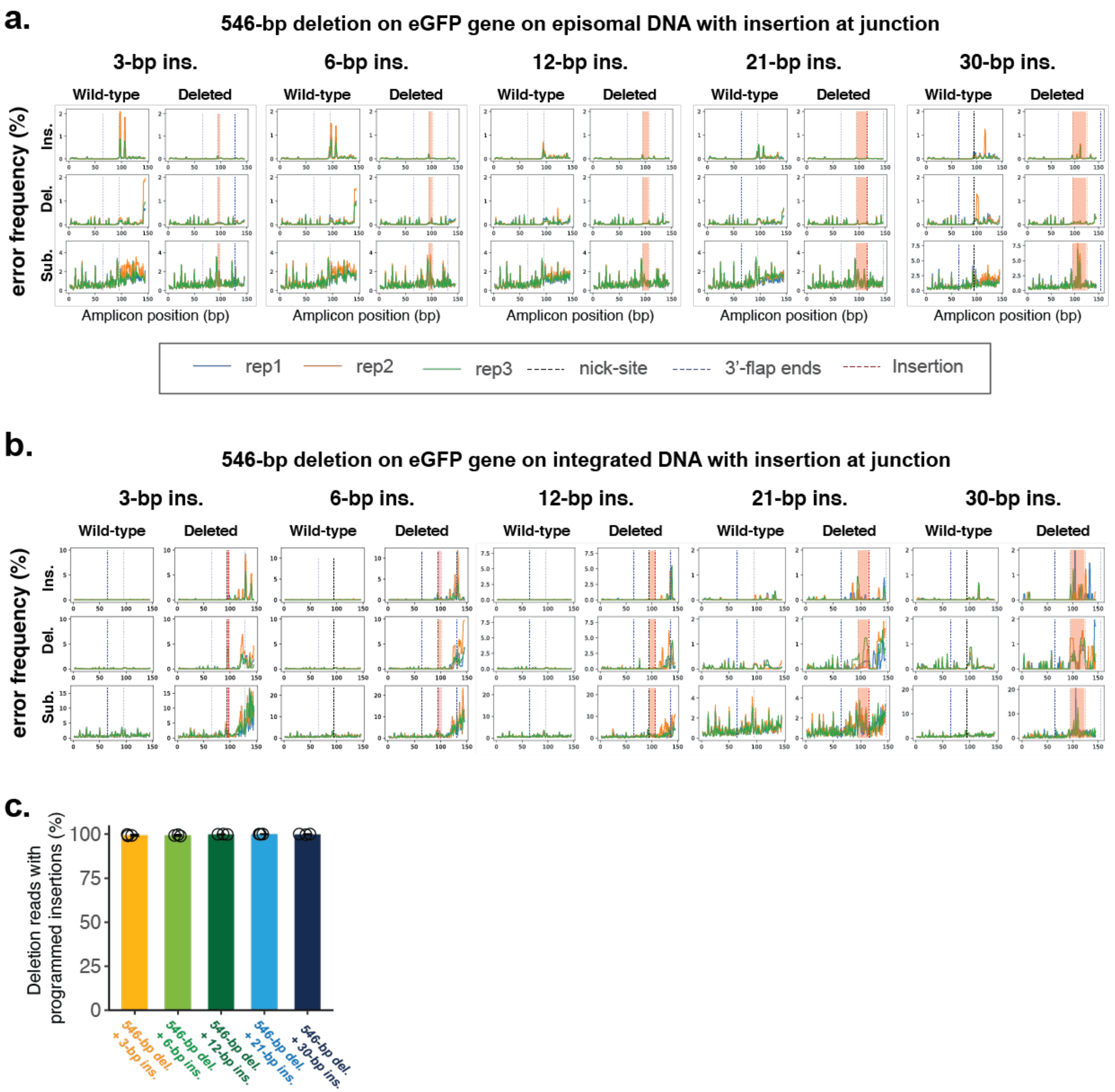


**Supplementary Figure 2. Error profiles with concurrent deletion and insertion at episomally or genomically encoded *eGFP*. a.** Insertion, deletion and substitution error frequencies plotted across sequencing reads from concurrent 546-bp deletion and various insertion conditions, targeting episomally encoded *eGFP*. These are based on single-end sequencing, with three replicates per experiment, all sequenced on one run, overlaid. Note that only one of the two 3'-DNA-flaps is covered by the sequencing read in amplicons lacking the deletion (labeled as 'wild-type'). Locations within read corresponding to insertions at deletion junction are highlighted between the nick-site (black dotted line) and end of insertion (red dotted line). *Y*-axis scaling is different for each plot. **b.** Same as (**a**), but for experiments targeting a genomically integrated copy of *eGFP*. **c.** The percentage of reads containing the programmed deletion that also contain the programmed insertion. Similar to **Fig. 2f**, but for experiments targeting a genomically integrated copy of *eGFP*. Error bars represent standard deviation for at least three replicates.

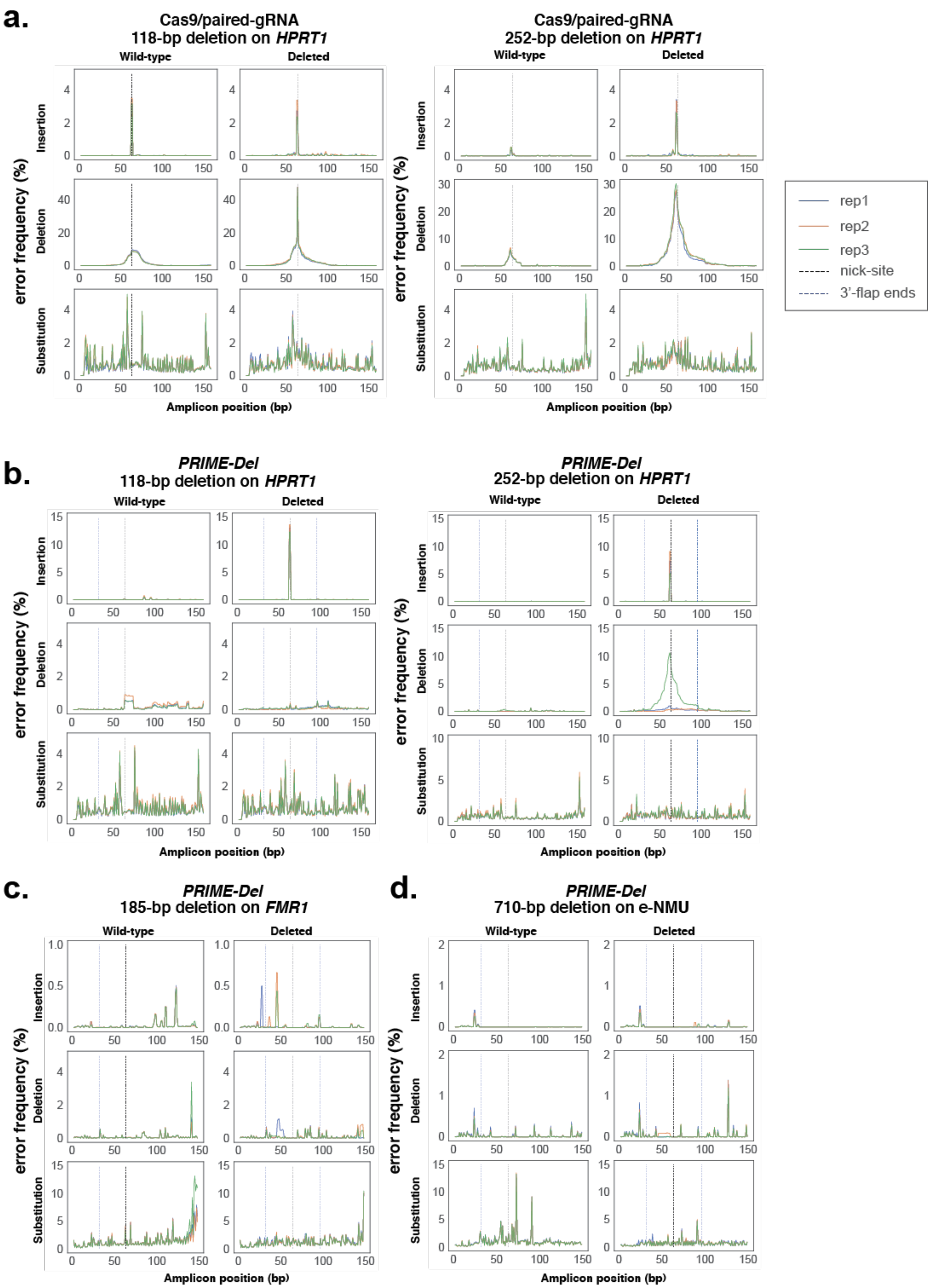
a.
Cas9/paired-gRNA
118-bp deletion on HPRT1
Cas9/paired-gRNA
252-bp deletion on HPRT1
b.
PRIME-Del
118-bp deletion on HPRT1
PRIME-Del
252-bp deletion on HPRT1
c.
PRIME-Del
185-bp deletion on FMR1
d.
PRIME-Del
710-bp deletion on e-NMU
Wild-type
Deleted
Insertion
Deletion
Substitution
error frequency (%)
Amplicon position (bp)
rep1
rep2
rep3
nick-site
3'-flap ends

**Supplementary Figure 3. Error profiles with targeting of native *HPRT1*, *FMR1*, and *e-NMU* loci. a-c.** Insertion, deletion and substitution error frequencies plotted across sequencing reads from: (**a**) 118-bp or 252-bp deletion on *HPRT1* using the Cas9/paired-gRNA strategy, (**b**) 118-bp or 252-bp deletion on *HPRT1* using the *PRIME-Del* strategy, **(c)** 185-bp deletion on *FMR1* using the *PRIME-Del* strategy, and (**d**) 710-bp deletion on *e-NMU* using the *PRIME-Del* strategy. Sequencing reads aligning to the 'deletion' reference for *HPRT1* condition are based on paired-end sequencing, while all the other conditions are based on the single-end sequencing. Each experiment has three replicates sequenced on one run, overlaid. Note that only one of the two 3'-DNA-flaps is covered by the sequencing read in amplicons lacking the deletion (labeled as 'wild-type') and that *y*-axis scaling is different for each plot. There may be a sample cross-contamination between Cas9/paired-gRNA replicate #1 and *PRIME-Del* replicate #3, based on the similarity of their deletion error profiles. Those two samples were PCR-amplified and AMPure processed in PCR tubes next to each other.

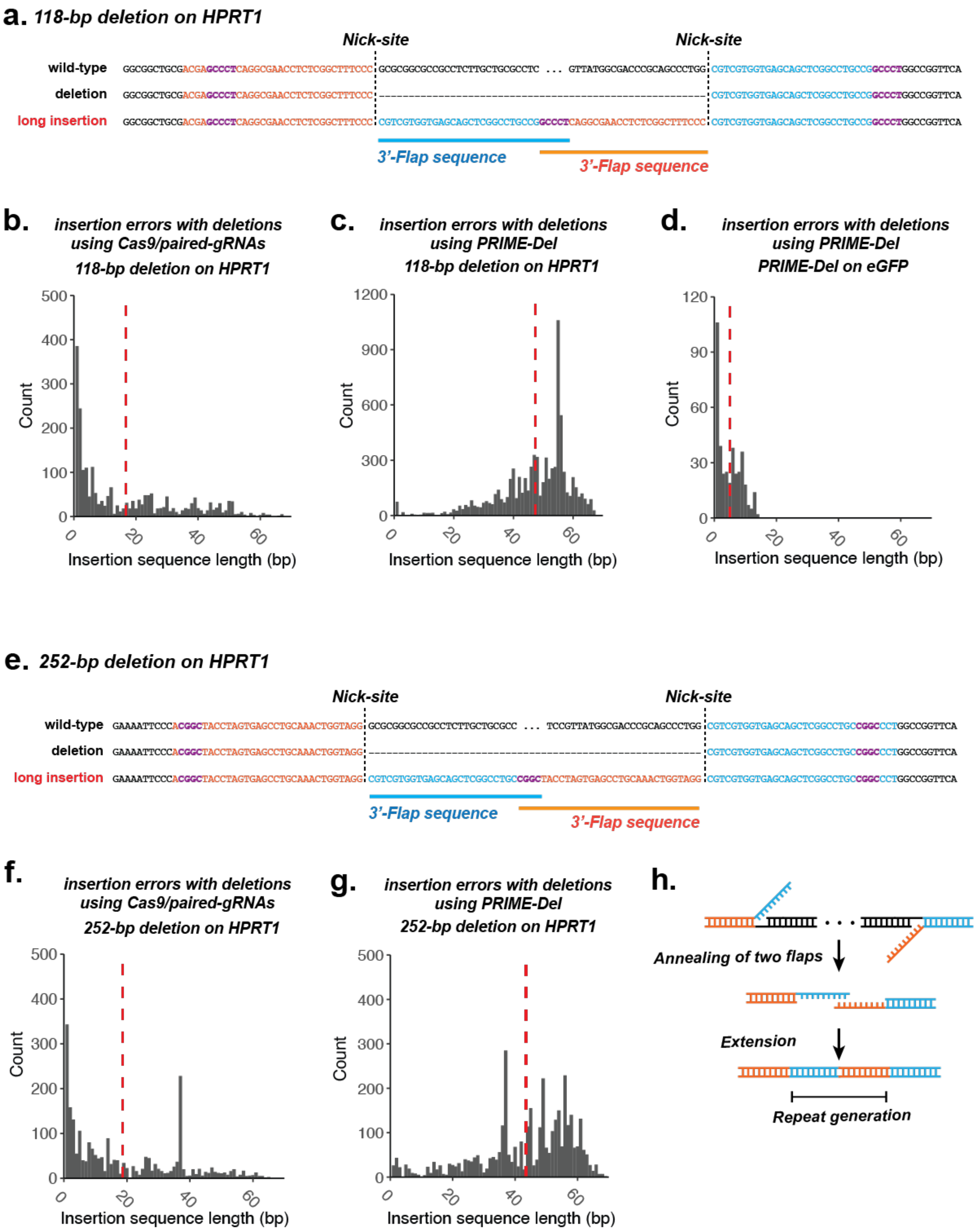
a. 118-bp deletion on HPRT1
Nick-site
Nick-site
wild-type GGCGGCTGCGACGAGCCCTCAGGCGAACCTCTCGGCTTTCCC GCGCGGCGCCGCCTCTTGCTGCGCCTC ... GTTATGGCGACCCGCAGCCCTGG CGTCGTGGTGAGCAGCTCGGCCTGCCGGCCCTGGCCGGTTCA
deletion GGCGGCTGCGACGAGCCCTCAGGCGAACCTCTCGGCTTTCCC CGTCGTGGTGAGCAGCTCGGCCTGCCGGCCCTGGCCGGTTCA
long insertion GGCGGCTGCGACGAGCCCTCAGGCGAACCTCTCGGCTTTCCC CGTCGTGGTGAGCAGCTCGGCCTGCCGGCCCTCAGGCGAACCTCTCGGCTTTCCC CGTCGTGGTGAGCAGCTCGGCCTGCCGGCCCTGGCCGGTTCA
3'-Flap sequence
3'-Flap sequence
b. insertion errors with deletions using Cas9/paired-gRNAs
118-bp deletion on HPRT1
Count
Insertion sequence length (bp)
c. insertion errors with deletions using PRIME-Del
118-bp deletion on HPRT1
Count
Insertion sequence length (bp)
d. insertion errors with deletions using PRIME-Del
PRIME-Del on eGFP
Count
Insertion sequence length (bp)
e. 252-bp deletion on HPRT1
Nick-site
Nick-site
wild-type GAAAATTCCCACGGCTACCTAGTGAGCCTGCAAACTGGTAGG GCGCGGCGCCGCCTCTTGCTGCGCC ... TCCGTTATGGCGACCCGCAGCCCTGG CGTCGTGGTGAGCAGCTCGGCCTGCCGGCCCTGGCCGGTTCA
deletion GAAAATTCCCACGGCTACCTAGTGAGCCTGCAAACTGGTAGG CGTCGTGGTGAGCAGCTCGGCCTGCCGGCCCTGGCCGGTTCA
long insertion GAAAATTCCCACGGCTACCTAGTGAGCCTGCAAACTGGTAGG CGTCGTGGTGAGCAGCTCGGCCTGCCGGCTACCTAGTGAGCCTGCAAACTGGTAGG CGTCGTGGTGAGCAGCTCGGCCTGCCGGCCCTGGCCGGTTCA
3'-Flap sequence
3'-Flap sequence
f. insertion errors with deletions using Cas9/paired-gRNAs
252-bp deletion on HPRT1
Count
Insertion sequence length (bp)
g. insertion errors with deletions using PRIME-Del
252-bp deletion on HPRT1
Count
Insertion sequence length (bp)
h.
Annealing of two flaps
Extension
Repeat generation

**Supplementary Figure 4. Rare long insertions upon *PRIME-Del* editing of the *HPRT1* exon 1. a.** We performed paired-end sequencing of amplicons derived from the *PRIME-Del*-edited *HPRT1* locus to bidirectionally cover the deletion junction and facilitate removal of PCR duplicates using 15-bp UMI sequences. This revealed recurrent long insertions that upon inspection appear to be chimeras of the two 3' flap sequences, with overlap at their GC-rich ends (highlighted in purple). Shown here is a representative insertion from the 118-bp deletion condition. **b-d.** Histograms of insertion sequence lengths for *HPRT1* 118-bp deletion with Cas9/paired-gRNA (**b**), *HPRT1* 118-bp deletion with *PRIME-Del* (**c**), or *eGFP* 546-bp deletion with *PRIME-Del* (**d**). Red vertical lines denote the mean insertion lengths. **e.** Same as (**a**), but representative insertion from the 252-bp deletion condition, also a chimera of the two 3' flap sequences, with overlap at their GC-rich ends. **f-g.** Histogram of insertion sequence lengths for *HPRT1* 252-bp deletion with *PRIME-Del* (**f**) or Cas9/paired-gRNA (**g**). **h.** Potential mechanism of long insertions with *PRIME-Del*. GC-rich ends of 3'-flaps of paired pegRNAs (*GCCCT* in case of 118-bp deletion and *CGGC* in case of 252-bp deletion) anneal to one another, or to another GC-rich stretch, resulting in insertion upon repair.

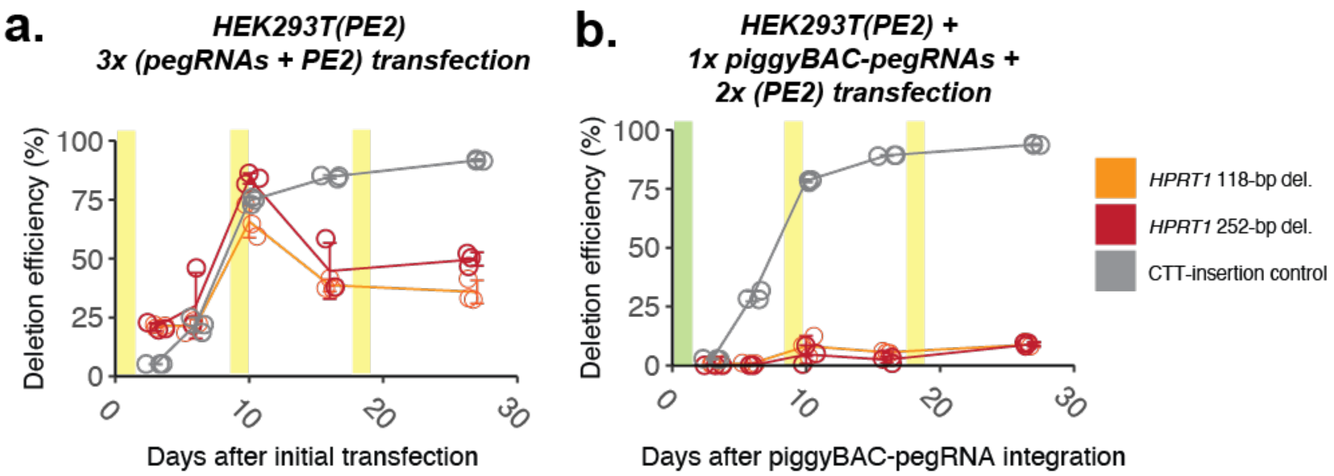


**Supplementary Figure 5. Multiple transfections enhance *PRIME-Del* efficiency in monoclonal HEK293T(PE2) cells. a.** Editing efficiencies measured for the 118-bp and 252-bp deletions at genomic *HPRT1* exon 1 using *PRIME-Del* (paired-pegRNA construct) or CTT-insertion using prime-editing (single-pegRNA construct), as a function of time after initial transduction of pegRNA(s). Plasmids bearing paired-pegRNAs and PE2 were transfected 3 times (days 0, 9, 18; highlighted in yellow) into PE2-expressing HEK293T cells. Error bars represent standard deviation for three replicates. **b.** Same as (**a**), but first with integration of pegRNAs to PE2-expressing HEK293T via piggyBAC transposon system on Day 0 (highlighted in green), followed by two additional transfections of plasmid bearing PE2 only on Day 9 and 18 (highlighted in yellow). Error bars represent standard deviation for three replicates.
